# Mosaic of local adaptation between white clover and rhizobia along an urbanization gradient

**DOI:** 10.1101/2023.08.16.553632

**Authors:** David Murray-Stoker, Marc T. J. Johnson

## Abstract

1. Urbanization frequently alters the biotic and abiotic environmental context. Variation in both the biotic and abiotic environment can result in selection mosaics that lead to locally adapted phenotypes. Ecological changes to urban environments have been found to result in evolutionary responses across a diverse array of plants and animals, but the effects of urbanization on local adaptation and coevolution between species remains largely unstudied.
2. Using a common garden experiment of 30 populations and 1080 plants, we tested for local adaptation in the mutualism between white clover (*Trifolium repens*) and rhizobia (*Rhizobium leguminosarum* symbiovar *trifolii*) along an urbanization gradient. We asked: (Q1) are white clover and rhizobia locally adapted? (Q2) Does nitrogen (N) addition mediate the strength of local adaptation? (Q3) Is the strength of local adaptation in host plants related to the strength of local adaptation in rhizobia, as expected if white clover and rhizobia are co-adapting? And (Q4) How does the degree of local adaptation vary with urbanization?
3. We found evidence for local adaptation for both white clover and rhizobia, with stronger signatures of local adaptation for rhizobia than white clover (Q1). While N addition positively affected plant biomass and negatively affected nodule density, N addition did not consistently mediate patterns of local adaptation (Q2). The strength of local adaptation was positively related between white clover and rhizobia under N addition, suggesting that soil N mediates co-adaptation between white clover and rhizobia (Q3). We did not find a consistent relationship between measures of urbanization and local adaptation (Q4). Additionally, fitness responses for white clover and rhizobia were unrelated to *Rhizobium* abundance and local adaptation was unrelated to the broader bacterial microbiome in the soil inoculants or root endosphere.
4. *Synthesis*. Our results show a spatial mosaic in local adaptation, with stronger local adaptation for rhizobia than white clover. While urbanization does influence the ecology of plant-microbe interactions, our study suggests urbanization does not disrupt local adaptation and coevolution among mutualists.

## Introduction

Urbanization is a dominant driver of environmental change that frequently alters the ecological setting of ecosystems (Pickett et al., 2001; Santangelo et al., 2022). Abiotic changes to the environment commonly include increased impervious surface, habitat fragmentation and degradation, elevated temperature, higher pollution (e.g., air, water, light, noise), and increased nutrient deposition (Grimm et al., 2008; Pickett et al., 2011). In addition to urban-driven abiotic changes, biotic changes frequently include altered species diversity and community composition, reduced vegetation cover, increased abundance of non-native species, and reduced abundance and diversity of native species (Faeth et al., 2011; McKinney, 2008). Ecological changes to urban environments have been found to cause evolution in a diverse array of plants and animals (Diamond & Martin, 2021; Johnson & Munshi-South, 2017). Despite increasing evidence of evolutionary consequences in response to urbanization, it remains to be seen if urbanization affects the local adaptation and (co)evolution of species interactions.

Variation in both the biotic and abiotic environment can result in selection mosaics that lead to locally adapted phenotypes (Thompson, 2005). Numerous environmental factors can impose natural selection that lead to local adaptation, whereby local populations have greater fitness than nonlocal populations (Blanquart et al., 2013). Mutualisms are a useful focal interaction to study local adaptation because the reciprocal fitness feedbacks among interacting species should lead to fitness alignment in a given environment but still allow for variation across environments (Heath & Stinchcombe, 2014; Hoeksema, 2010; Nuismer et al., 1999).

Variation in community composition across space and time can alter mutualistic species interactions (Thompson, 1988, 2005). Abiotic context can also affect mutualisms by altering the physical (e.g., temperature; Kiers *et al*. 2010) or resource (e.g., nutrient and water availability; Bronstein 1994; Shantz *et al*. 2016) environment in which species interact. Moreover, biotic and abiotic contexts are likely acting in concert to affect the outcome of mutualistic interactions (Burghardt et al., 2022; Heath et al., 2010; Heath & Tiffin, 2007; Van Cauwenberghe et al., 2016). Because urbanization alters both the biotic and abiotic environment simultaneously across spatial and temporal scales (Diamond & Martin, 2021; Johnson & Munshi-South, 2017), urban environments have the potential to alter selective landscapes, local adaptation, and potentially coevolution between interacting species.

Mutualistic interactions between legumes and rhizobia are a useful model for evaluating evolutionary ecology in urban environments. In this mutualism, rhizobia fix atmospheric N_2_ into an accessible form of nitrogen (N) for the plant (NH ^+^) in exchange for photosynthate and hosting within nodules provided by the plant (Poole et al., 2018). Ecological changes in urban environments, particularly increased N deposition and enrichment (Grimm et al., 2008; Stevens et al., 2018), could have consequences for legume-rhizobium local adaptation by disrupting the economy of the resource exchange: direct acquisition of N is ‘cheaper’ for the host legume, which can select for less cooperative rhizobia (Regus et al., 2017; Weese et al., 2015). In a previous study along an urbanization gradient (MurrayLStoker & Johnson, 2021), we found that urbanization altered the ecology of a legume-rhizobia mutualism. Specifically, urban plants exhibited reduced investment in the mutualism and primarily acquired N through fixation except at the urban and rural limits of the gradient. Soil N content was the nexus linking the effects of urbanization to the mutualism (MurrayLStoker & Johnson, 2021). With this evidence, we wanted to determine if the ecological changes driven by urbanization affected local adaptation of white clover and its rhizobia mutualists.

Here we conducted a large, factorial field experiment to test the hypothesis that urban environmental change drives local adaptation in the mutualism between white clover (*Trifolium repens*) and rhizobia (*Rhizobium leguminosarum* symbiovar *trifolii*). We asked four specific questions: **(Q1)** Are white clover and rhizobia locally adapted? **(Q2)** Does N addition mediate the strength of local adaptation? **(Q3)** Is the strength of local adaptation in host plants related to the strength of local adaptation in rhizobia, as expected if white clover and rhizobia are coadapting? **(Q4)** How does the degree of local adaptation vary with urbanization? Results from this study provide the first test of how urbanization affects the evolution and local adaptation of interacting species, which provides insight into urban environments and whether they could be disrupting coevolution.

## Materials and Methods

### Study System

We used the mutualism between white clover (*Trifolium repens* L., Fabaceae) and its facultative, rhizobial symbiont (*Rhizobium leguminosarum* symbiovar *trifolii*) to evaluate if and how local adaptation varied with urbanization. White clover is a perennial, herbaceous legume native to Eurasia that is now globally distributed. White clover typically reproduces clonally through stolons as well as through seed via obligately-outcrossed flowers (Burdon, 1983). *Rhizobium leguminosarum* symbiovar *trifolii* (hereafter rhizobia) is the primary rhizobial symbiont of white clover (Andrews & Andrews, 2017).

### Biological Materials

*White clover:* The 30 focal white clover populations for this experiment were a subset of those sampled by Murray-Stoker and Johnson (2021), with focal populations selected to span an urbanization gradient in the Greater Toronto Area, Ontario, Canada (Figure 1A). To minimize parental effects, we grew field-collected seeds in the greenhouse to produce outcrossed F_1_ populations. All F_1_ seeds were derived from a random panmictic cross among 5-10 parent plants within a source population (detailed methods are provided in Appendix S1).

**Figure 1:**
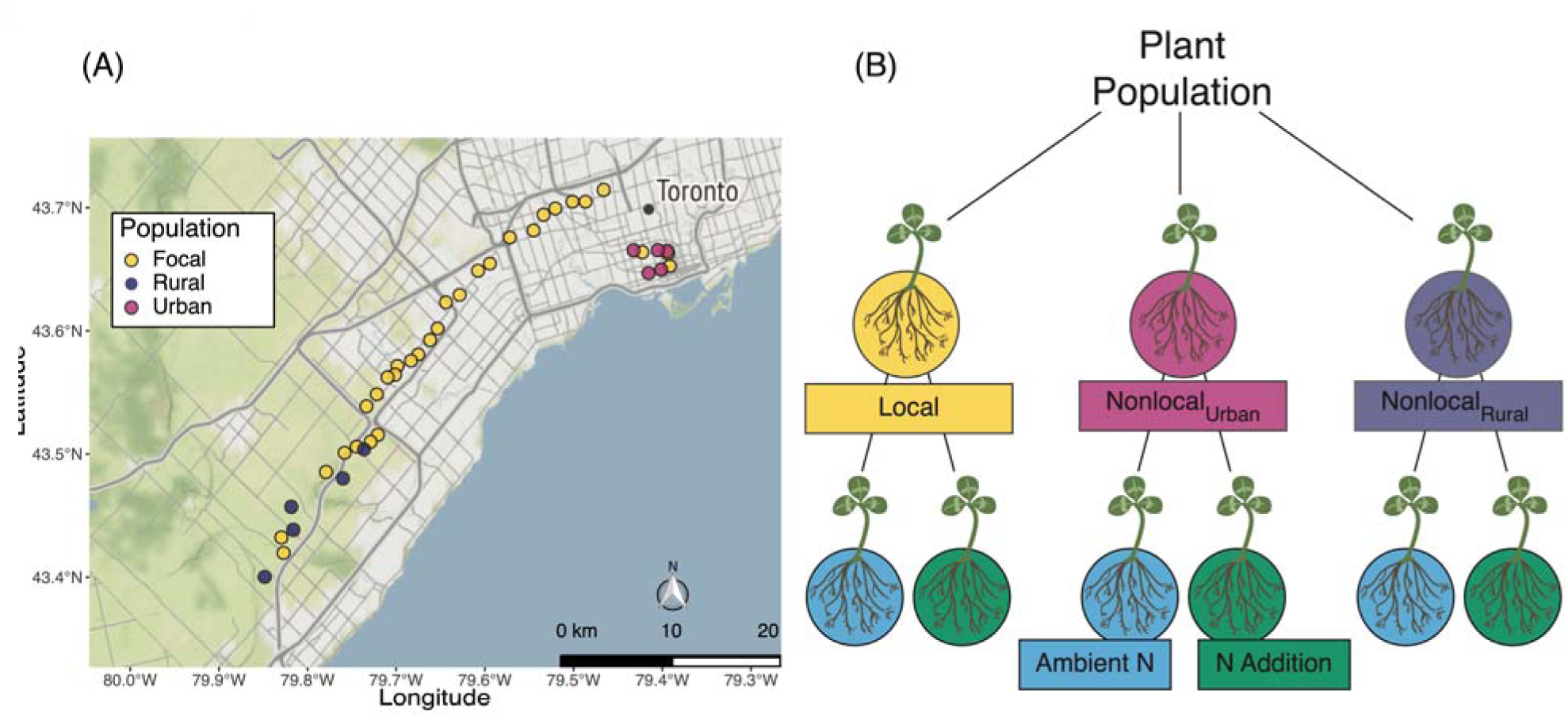
Map of the urbanization gradient in the Greater Toronto Area, Ontario, Canada (A), with the location of the 30 focal populations indicated by yellow circles. We also show the location of the 5 rural (blue circles) and 5 urban (pink circles) sites from which nonlocal inoculants were sourced. Urban and suburban areas in gray, nonurban agricultural and forested areas in green, and Lake Ontario in light blue. Map tiles by Stamen Design, under CC BY 3.0. Data by OpenStreetMap, under ODbL. We also show a diagram of the experimental design (B). Each of the 30 focal populations (yellow circles on the map) received 3 microbiome treatments (local, nonlocal_Urban_, and nonlocal_Rural_). Additionally, each microbiome treatment received 2 nitrogen treatments (no addition = ambient N, 10 mM KNO_3_ fertilizer = N addition). The factorial experiment was conducted in an outdoor common garden at the University of Toronto Mississauga campus.

*Soil inoculants:* Soil for microbiome inoculants was collected following the methods described in Murray-Stoker and Johnson (2021). Briefly, 5 soil cores were taken at each site at least 5 m from the nearest white clover plant and other legumes (e.g., *Medicago lupulina*). Each soil core was collected to a depth of 5 cm, with the top 2 cm plus any organic material removed. Individual cores were stored in separate ziploc bags and stored at −80°C until subsequent processing. Local microbiomes were paired to each of the 30 white clover populations in the experiment, while the nonlocal_Urban_ and nonlocal_Rural_ microbiome inoculants were pooled from 5 populations at the urban or rural ends of the gradient, respectively (Figure 1A).

We used a local versus nonlocal microbiome inoculation approach (Blanquart et al., 2013; Heath & Stinchcombe, 2014) to test for local adaptation between white clover and rhizobia for two principal reasons. First, this approach tests for overall differences in fitness between local and nonlocal white clover or rhizobia populations, which can be further distinguished based on the habitat of populations (i.e., urban vs. rural). In particular, increased N deposition in urban environments can alter the dynamics of the mutualism between white clover and rhizobia by reducing nodule densities and influencing the source of N used by white clover (MurrayLStoker & Johnson, 2021). Additionally, urbanization leads to homogenization of the biotic and abiotic environment (McKinney, 2006; Santangelo et al., 2022), which might result in homogenized composition of microbial communities (Delgado-Baquerizo *et al*. 2021; Rosier *et al*. 2021; but see Epp Schmidt *et al*. 2017). Together, this evidence suggests that the degree of local adaptation might depend on (1) the nonlocal inoculant representing rural or urban environments and (2) similarity in microbiome composition between the local and either nonlocal microbiomes (i.e., nonlocal_Rural_ and nonlocal_Urban_). Second, by using microbiome inoculants instead of isolated rhizobia strains, this approach also provided more ecological realism by allowing rhizobia and other bacteria and fungi to establish in the soil and roots.

### Experiment Overview

We tested for local adaptation between populations of white clover and rhizobia using a common garden experiment (Figure 1B). We used the local versus nonlocal contrast (sensu sympatric vs. allopatric; Blanquart *et al*. 2013), with local adaptation indicated when mean local fitness was greater than mean nonlocal fitness. We crossed 30 white clover populations with 3 microbiome treatments (local, nonlocal_Rural_, and nonlocal_Urban_) and 2 N treatments (no N addition = ambient N, 10 mM KNO_3_ fertilizer = N addition). Each treatment combination had 6 replicates per population (30 populations × 3 microbiome treatments × 2 N treatments × 6 replicates = 1080 plants). The N addition treatment was representative of the soil N content measured at the focal populations by Murray-Stoker and Johnson (2021) and was designed to reflect the impacts of urbanization on soil N. We also had 6 control pots (3 ambient N, 3 N addition) that were not planted or inoculated but did receive fertilizer additions, and these pots were used to evaluate dispersal and establishment of bacteria at the experimental site.

### Common Garden Experiment

We grew all of the plants outside in a common environment using an incomplete block design at the University of Toronto Mississauga campus. In total, there were 9 blocks consisting of 120 plants; each block included all treatment combinations but not all population-by-treatment combinations were represented in each block due to spatial constraints. Prior to planting, seeds were scarified with sandpaper, surface sterilized (60 s in 70% ethanol, 60 s in 10% household bleach, 90 s rinse in sterile water) and then germinated in the dark on dampened filter paper for 48 h at 20°C. Germinated seeds were planted in 1 L bleach-sterilized pots in a steam-sterilized (2 h of sterilization) mix of potting soil and sand mixture (2:3 V/V). The soil mixture was selected to facilitate harvesting of all root biomass at the end of the experiment. We planted two germinated seeds in each pot, and pots were thinned to one plant between weeks 2-3 of the experiment.

Microbiome treatments were applied 1 week after planting by applying soil slurries. Soil slurries for each microbiome treatment were prepared with 7.5 g fresh soil (thawed from –80°C overnight at 4°C) in 50 mL sterile deionized water; nonlocal treatments were composed of equal mixtures by mass from 5 populations within the respective habitat type. Slurries were hand shaken for 60 s after preparation to mix and suspend the soil, with further re-suspension (hand shaken for 30 s) during inoculation to maintain soil suspension. Each plant received 10 mL of its respective microbiome inoculant. Weekly N addition treatments began in week 2, and we applied bi-weekly N-free fertilizer following Batstone *et al*. (2017) starting in week 4 of the experiment. Both the N-addition and N-free fertilizers were autoclave sterilized (60 min sterilization) prior to application. Plants were watered daily as needed for the duration of the experiment. We ran the experiment for 19 weeks (21 May−2 October 2021). Due to germination failure, herbivory, and other sources of plant loss, our final plant count was 841.

### Laboratory Processing

We measured multiple components of fitness for white clover (aboveground biomass and belowground biomass) and rhizobia (total nodule density and fixing nodule density). We first washed all plants of soil and debris and then split them into aboveground and belowground components to facilitate fitness measurements. Aboveground components were placed in envelopes, dried to constant mass (at least 72 h at 60°C), and aboveground biomass was weighed to the nearest 1 mg. To measure rhizobia fitness components, a subset of roots was collected from each plant by cutting the first three roots below the plant base and directly attached to the stolon (following MurrayLStoker & Johnson 2021), and we then measured the total root length of all collected roots (mean ± standard error = 43.5 ± 0.58 cm, range = 2.1-128.7 cm). Nodule density, an estimate of investment in the mutualism by white clover and rhizobia fitness, was measured by visually-counting nodules from each measured root and standardizing by the total root length to generate an estimate of nodule density per cm of root per plant (MurrayLStoker & Johnson, 2021). We also calculated the density of fixing nodules, a measure of the effectiveness of the mutualism, by counting the number of nodules with pink colouration (indicative of N fixation; Udvardi & Poole 2013) and calculating density as described above. After estimating nodule density and fixing nodule density, all belowground biomass (including the 3 excised roots) was placed in an envelope, dried for 72 h at 60°C, and weighed to the nearest 0.1 mg. We focused on biomass as a measure of plant fitness because white clover undergoes extensive clonal growth as a perennial. We could not measure sexual fitness accurately because only a fraction of plants flowered and we could not prevent interpopulation crosses.

### DNA Extraction & Sequencing

To determine if white clover and rhizobia fitness responses were influenced by the broader bacterial microbiome, we sequenced the bacterial microbiome for each of the microbiome inoculants and a subset of roots from each of the experimental treatments. All 40 microbiome inoculants (30 focal, 5 nonlocal_Rural_, 5 nonlocal_Urban_) were sequenced to evaluate variation in bacterial community composition. A total of 60 root microbiomes (10 populations × 3 microbiome treatments × 2 N treatments = 60 plants) were sequenced to evaluate: (1) white clover and rhizobia fitness responses as a function of *Rhizobium* abundance, (2) *Rhizobium* abundance in response to experimental treatments, and (3) fitness of white clover and rhizobia in response to soil and root bacterial microbiome composition. We included each treatment combination for the same 6 populations (i.e., 10 populations × 6 treatment combinations = 60 samples). Protocols for microbiome DNA extraction are provided in the supplementary material (Appendix S1).

### Statistical Analyses

We provide detailed methods for all statistical analyses in the supplementary material (Appendix S1). Briefly, we analyzed variation in fitness responses for white clover (aboveground biomass, belowground biomass) and rhizobia (nodule density, fixing nodule density) in response to the microbiome treatment (local, nonlocal_Rural_, nonlocal_Urban_), N treatment (ambient N, N addition), and the two-way interaction using linear mixed-effects models with Type III sums-of-squares.

We also quantified the extent to which population × environment interactions were due to changes in population variance and changes in rank order using Cockerhams’s method (Cockerham, 1963; Muir et al., 1992). To test if there was a general signature of local adaptation, we repeated the same procedure described above but with microbiome recoded as local and nonlocal_Global_ (i.e., nonlocal_Rural_ and nonlocal_Urban_ combined).

The strength of local adaptation was calculated using three indices: (1) local − nonlocal_Global_, (2) local − nonlocal_Rural_, and (3) local − nonlocal_Urban._ These indices were calculated separately for each of the four fitness response variables (i.e., aboveground biomass, belowground biomass, nodule density, and fixing nodule density). Positive values were indicative of local adaptation because they showed greater fitness in the local versus nonlocal treatment. Local adaptation indices were calculated using the best linear unbiased predictors (i.e., BLUPs) from linear mixed-effects models. To test whether the strength of local adaptation was related between interacting white clover and *Rhizobium* populations (i.e., a signature of coadaptation), we calculated all pairwise Pearson correlations between the two fitness components of white clover with the two fitness components of rhizobia. We calculated correlations separately for each N treatment to examine if N addition mediated coadaptation.

To test how urbanization affects the strength of local adaptation, we regressed local adaptation indices against measures of urbanization: distance from the city center, mean impervious surface cover (ISC, 250 m radius around each population), and the Human Influence Index (HII, 1 km radius around each population). We fitted separate regressions for each local adaptation index and urbanization metric due to multicollinearity between variables (variance inflation factors > 5–10). We scaled all predictor variables to a mean of 0 and standard deviation of 1 prior to analysis to allow for direct comparisons across models.

Microbiome sequences were processed for analysis using the DADA2 pipeline (Callahan et al., 2016), with taxonomy assigned to amplicon sequence variants using the Ribosomal Database Project (Cole et al., 2014) and naïve Bayesian classifier (Wang et al., 2007) implemented in DADA2. We regressed white clover and rhizobia fitness responses against *Rhizobium* abundance (number of reads assigned to the genus *Rhizobium*), N treatment, and the interaction using generalized linear models. We examined differences in *Rhizobium* abundance in response to microbiome_Global_ (local vs. nonlocal_Global)_, N (ambient N, N addition), and the interaction using ANOVAs with Type III sums-of-squares. Finally, we examined the role of microbiome composition on local adaptation. We tested if the strength of local adaptation increased when microbiome inoculant composition was more dissimilar (pairwise Bray-Curtis dissimilarity) using a linear regression. We also tested if local adaptation indices were related to root bacterial composition. The effects of the microbiome treatment, N treatment, and their interaction on root microbiome composition was fit and assessed using PERMANOVA with a Bray-Curtis dissimilarity matrix (Oksanen et al., 2020). Relationships between root community composition and local adaptation indices were quantified using the ’envfit()’ function (Oksanen et al., 2020).

All statistical analyses were performed using R (version 4.2.2; R Core Team 2022) in the RStudio environment (version 2022.07.2; RStudio Team 2022). All packages used for data management, analysis, and figure creation are provided in the detailed statistical analyses in the supplementary material (Appendix S1). We followed statistical best practices by focusing on estimates, variation of the estimates, and effect sizes to interpret the results and identify the strength of evidence (Berner & Amrhein, 2022; Gelman & Hennig, 2017; McShane et al., 2019; Wasserstein et al., 2019; Wasserstein & Lazar, 2016).

## Results

### Effects of microbiome and nitrogen treatments on fitness

We found evidence for local adaptation for both white clover and rhizobia, and the effects were stronger for rhizobia than white clover and varied among populations (Table 1, Figure 2). White clover demonstrated weak adaptation to local microbiomes for both aboveground biomass (P = 0.047, ^2^ = 0.106, Table 1, Figure 2) and belowground biomass (P = 0.036, ^2^ = 0.008, Table 1, Figure 2). Aboveground biomass was 6.4% and 12.8% greater in local versus nonlocal_Rural_ and nonlocal_Urban_ microbiomes, respectively (contrasts in Table S1), and 60% of populations demonstrated patterns consistent with local adaptation (Figure 2). In contrast, belowground biomass was nearly equivalent (0.8% increase) between local and nonlocal_Rural_ microbiomes, whereas belowground biomass was 7.4% greater in local than nonlocal_Urban_ microbiomes (contrasts in Table S1).

**Figure 2:**
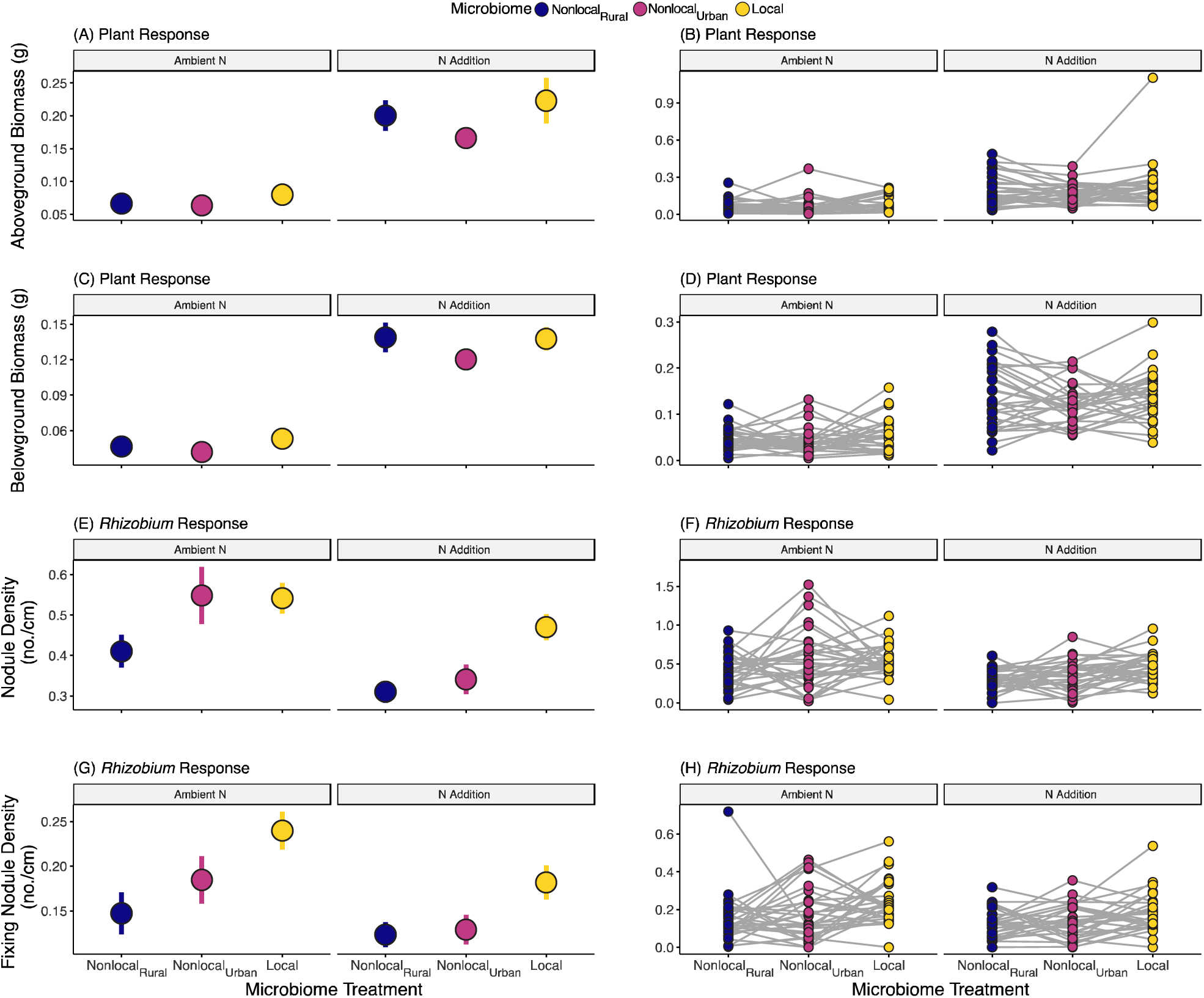
Treatment means ± 1 SE (left column) and reaction norms of population means (right column) by microbiome and nitrogen treatments for aboveground biomass (A-B), belowground biomass (C-D), nodule density (E-F), and fixing nodule density (G-H). Microbiome treatments are indicated as nonlocal_Rural_ (blue), nonlocal_Urban_ (pink), and local (yellow); individual plots are faceted by nitrogen treatment (ambient N and N addition). Inset text indicates the factors of interest in the optimized model structure for the respective variable, with fixed effects in regular font and random effects in italics. Influential factors and interactions were identified based on effect sizes (η^2^_p_ range = 0.106-0.871, ICC range = 0.035-0.126) and are in bold, with detailed test statistics provided in Table 1.

**Table 1:** Summary of the linear mixed-effects models testing the effects of microbiome (local, nonlocal_Rural_, nonlocal_Urban_), nitrogen (ambient N, N addition), and the two-way interaction on measure of plant and rhizobia fitness. We provide the numerator degrees-of-freedom (NumDF), denominator degrees-of-freedom (estimated using the Kenward-Roger method (DenDF; Kenward & Roger 1997), F-statistics calculated from type III sums-of-squares, P-values, and the effect size (partial eta-squared, χ^2^_P_) for each fixed effect term in the ANOVAs. We also provide the *Χ*^2^ statistic, P-values from likelihood ratio tests, and the effect size (intra-class correlation coefficient, ICC) for each random effect term, with random effect terms and statistics in italics. The variance explained by the fixed effects (R^2^_Marginal_) and combined fixed and random effects (R^2^_Conditional_) is provided for each model.

| Term | NumDF | DenDF | F/$\chi^2$ | P-value | $\eta^2_p$/ICC |
| --- | --- | --- | --- | --- | --- |
| **Aboveground Biomass** |  |  |  |  |  |
| **($R^2_{\text{Marginal}} = 0.233$, $R^2_{\text{Conditional}} = 0.397$)** |  |  |  |  |  |
| Microbiome | 2 | 54.911 | 3.243 | 0.047 | 0.106 |
| Nitrogen | 1 | 27.790 | 188.399 | < 0.001 | 0.871 |
| Microbiome × Nitrogen | 2 | 742.517 | 0.470 | 0.625 | 0.001 |
| *Microbiome × Population* |  |  | 4.951 | 0.026 | 0.040 |
| *Nitrogen × Population* |  |  | 3.901 | 0.048 | 0.035 |
| *Population* |  |  | 10.311 | 0.001 | 0.126 |
| *Block* |  |  | 3.575 | 0.059 | 0.011 |
| **Belowground Biomass** |  |  |  |  |  |
| **($R^2_{\text{Marginal}} = 0.292$, $R^2_{\text{Conditional}} = 0.376$)** |  |  |  |  |  |
| Microbiome | 2 | 802.275 | 3.338 | 0.036 | 0.008 |
| Nitrogen | 1 | 801.735 | 384.570 | < 0.001 | 0.324 |
| Microbiome × Nitrogen | 2 | 805.884 | 0.292 | 0.747 | 0.001 |
| *Population* |  |  | 46.239 | < 0.001 | 0.104 |
| *Block* |  |  | 4.791 | 0.029 | 0.015 |
| **Nodule Density** |  |  |  |  |  |
| **($R^2_{\text{Marginal}} = 0.061$, $R^2_{\text{Conditional}} = 0.261$)** |  |  |  |  |  |
| Microbiome | 2 | 56.305 | 9.257 | < 0.001 | 0.247 |
| Nitrogen | 1 | 27.932 | 13.328 | 0.001 | 0.323 |
| Microbiome × Nitrogen | 2 | 55.584 | 1.057 | 0.354 | 0.037 |
| *Microbiome × Population* |  |  | 1.614 | 0.204 | 0.053 |
| *Nitrogen × Population* |  |  | 0.019 | 0.889 | 0.004 |
| *Microbiome × Nitrogen × Population* |  |  | 6.376 | 0.012 | 0.105 |
| *Population* |  |  | 1.899 | 0.168 | 0.047 |
| *Block* |  |  | 0.390 | 0.533 | 0.003 |
| **Fixing Nodule Density** |  |  |  |  |  |
| **($R^2_{\text{Marginal}} = 0.043$, $R^2_{\text{Conditional}} = 0.198$)** |  |  |  |  |  |
| Microbiome | 2 | 55.390 | 10.067 | < 0.001 | 0.267 |
| Nitrogen | 1 | 27.929 | 5.595 | 0.025 | 0.167 |
| Microbiome × Nitrogen | 2 | 741.058 | 0.442 | 0.643 | 0.001 |
| *Microbiome × Population* |  |  | 8.159 | 0.004 | 0.062 |
| *Nitrogen × Population* |  |  | 5.201 | 0.023 | 0.045 |
| *Population* |  |  | 0.787 | 0.375 | 0.026 |
| *Block* |  |  | 13.511 | < 0.001 | 0.029 |

Signatures of local adaptation were stronger for rhizobia when estimated as either nodule density (P < 0.001, ^2^ = 0.247) or fixing nodule density (P < 0.001, ^2^ = 0.267, Table 1, Figure 2). Nodule density was 38.4% and 20.1% greater in local versus nonlocal_Rural_ and nonlocal_Urban_ microbiomes, respectively; however, nodule density depended on the three-way interaction between microbiome, N, and population (Table 1, Figure 2). There was an increase in the frequency of populations demonstrating local adaptation under N addition (local vs. nonlocal_Rural_ = 83%, local vs. nonlocal_Urban_ = 62%) compared to ambient N (local vs. nonlocal_Rural_ = 63%, local vs. nonlocal_Urban_ = 57%; contrasts in Table S1). Fixing nodule density was 53.4% and 33.8% greater in the local microbiome treatment than the nonlocal_Rural_ and nonlocal_Urban_ treatments, respectively (contrasts in Table S1).

As expected, N addition had strong and positive effects on aboveground biomass (81.3% increase, P < 0.001, η^2^_P_ = 0.871, Table 1, Figure 2) and belowground biomass (75.0% increase, P < 0.001, η^2^_P_ = 0.324, Table 1, Figure 2;contrasts in Table S1). In contrast, N addition decreased nodule density by 17.9% for both total nodule fixing density (P < 0.001, ^2^ = 0.323, Table 1, Figure 2) and fixing nodule density (P = 0.025, η2P = 0.167, Table 1, Figure 2; contrasts in Table S1). Notably, differences between local and nonlocal microbiomes were greater under N addition for both nodule density and fixing nodule density (Figure 1). When we further evaluated microbiome × population interactions using Cockerham’s method, we found that crossing of reaction norms (i.e., changes in rank order of populations among environments) accounted for most of the variation in the interaction for aboveground biomass (97.4%), nodule density (86.9%), and fixing nodule density (97.9%), as opposed to changes in population variance. Results described above were similar when microbiome treatments were examined as local vs. nonlocal_Global_: there were stronger effects of local adaptation for both nodule density and fixing nodule density but no evidence of local adaptation for belowground biomass (Table S2, Figure S2).

### Reciprocal Local Adaptation

White clover and rhizobia exhibited patterns of reciprocal local adaptation consistent with coevolution, but linkages between white clover and rhizobia depended on the N environment and the local – nonlocal contrast (Figures 3, S2-S3). A positive correlation between indices of local adaptation for aboveground biomass and nodule density was apparent under N addition for the global (local − nonlocal_Global_, r and 95% confidence interval: r = 0.418 [0.061, 0.681]; Figure 3), rural (local − nonlocal_Rural_, r = 0.396 [0.034, 0.666]; Figure S2), and urban contrasts (local − nonlocal_Urban_, r = 0.301 [−0.081, 0.606]; Figure S3). There was no clear correlation between indices of local adaptation for aboveground biomass and nodule density under ambient N (local − nonlocal_Global_, r = −0.194, 95% CI [−0.518, 0.179]; local − nonlocal_Rural_, r = −0.095, 95% CI [−0.440, 0.274]; local − nonlocal_Urban_, r = −0.087, 95% CI [−0.434, 0.282]; Figures 3, S2-S3). Relationships between belowground biomass and nodule density generally switched from negative under ambient N (local − nonlocal_Global_, r = −0.301 [−0.597, 0.067]; local − nonlocal_Rural_, r = −0.214 [−0.533, 0.159]; local − nonlocal_Urban_, r = −0.197 [−0.520, 0.176]; Figures 3, S2-S3) to positive under N addition (local − nonlocal_Global_, r = 0.253 [−0.125, 0.567]; local − nonlocal_Rural_, r = 0.305 [−0.069, 0.604]; local − nonlocal_Urban_, r = 0.052 [−0.327, 0.417]; Figures 3, S2-S3). When we examined the potential for reciprocal local adaptation between fixing nodule density and either aboveground or belowground biomass, we observed positive correlations in the rural contrast under N addition (aboveground biomass: local − nonlocal_Rural_, r = 0.424 [0.069, 0.684]; belowground biomass: local − nonlocal_Rural_, r = 0.399 [0.038, 0.668]; Figure S2).

**Figure 3:**
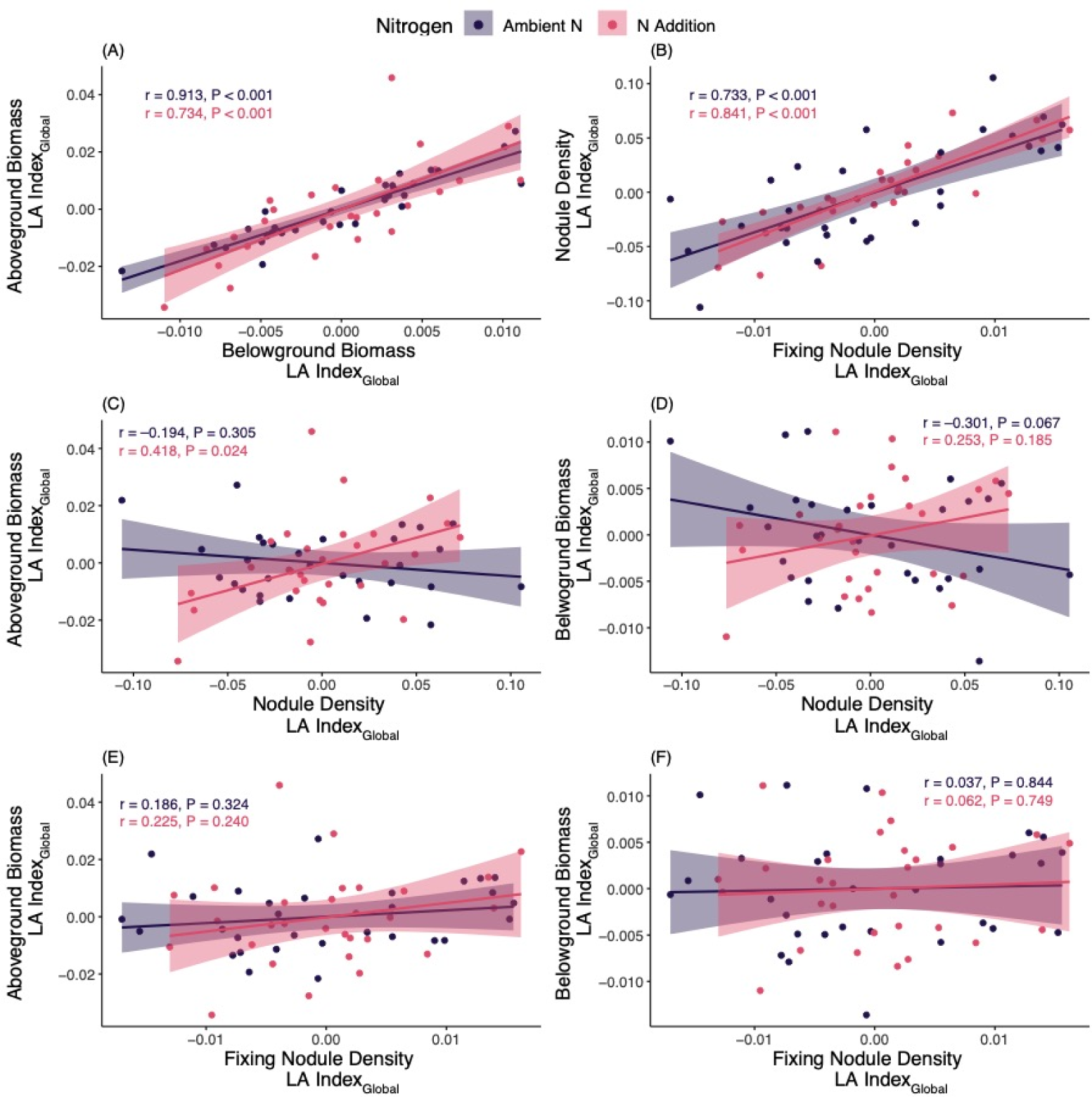
Pairwise correlations between the local – nonlocal_Global_ local adaptation indices for aboveground biomass, belowground biomass, nodule density, and fixing nodule density. Separate correlations were calculated for each nitrogen treatment (ambient N = purple, N addition = pink). Lines represent lines-of-best-fit (± 95% confidence interval). Inset text provides the correlation coefficient (r) and P-value for each correlation.

### Local Adaptation and Urbanization

We did not identify any strong or consistent relationships between local adaptation and urbanization, regardless of local adaptation index (Figures 4, S4-S6). There were no consistent patterns in the strength and direction of relationships between local adaptation indices for our fitness estimates and urbanization metrics (Figures 4, S4-S6), and all of the relationships explained a negligible amount of variation (range of R^2^ = 0-0.031).

**Figure 4:**
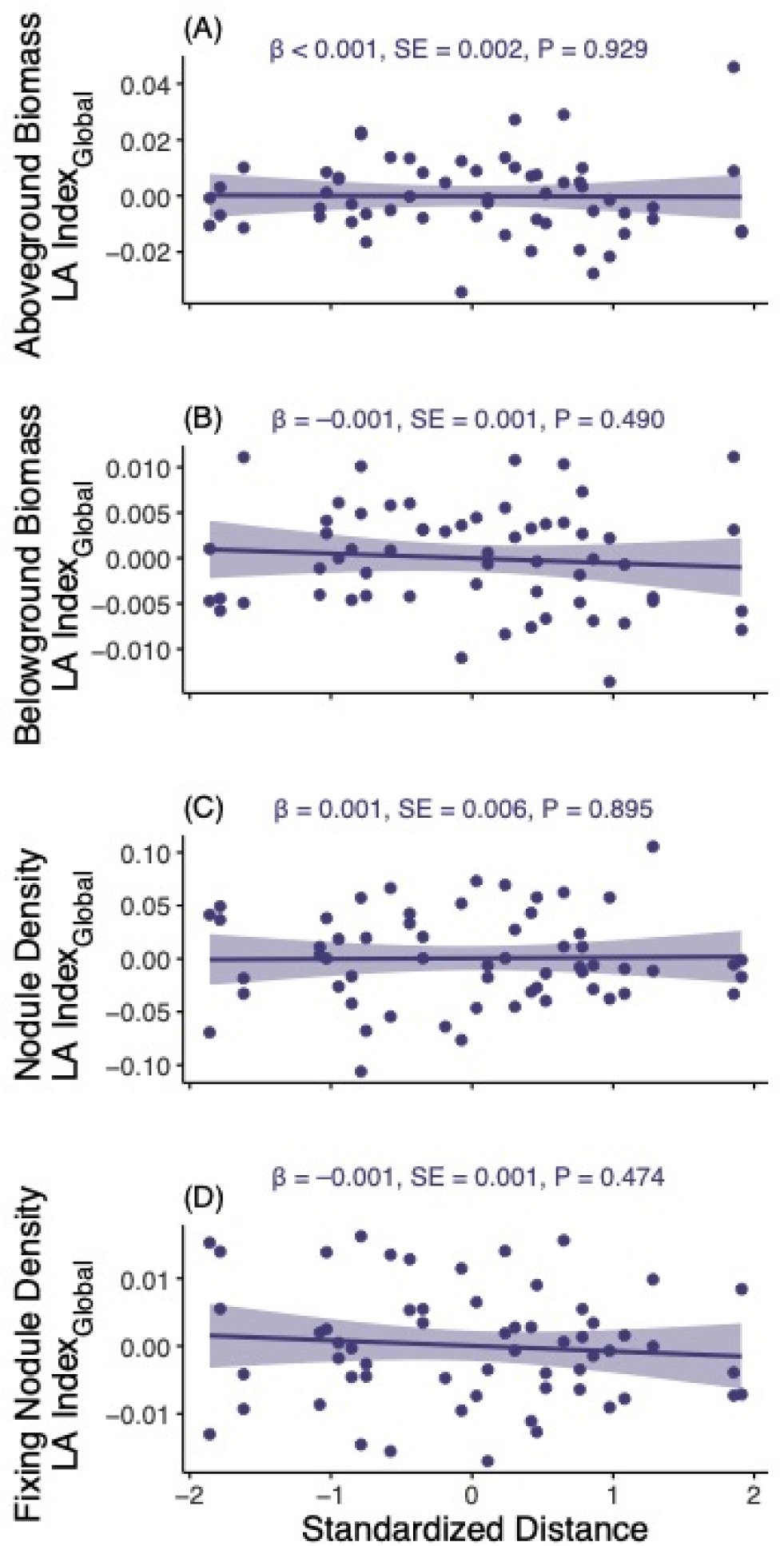
Relationship between the global local adaptation index (local − nonlocal_Global_, LA Index_Global_) and urbanization (standardized distance from the city centre; mean = 0, standard deviation = 1) for each of the fitness estimates [aboveground biomass (A) belowground biomass (B), nodule density (C), and fixing nodule density, (D)]. Lines represent the lines-of-best-fit (± 95% confidence interval). Inset text provides the slope parameter (β), standard error (SE), and P-value for each linear regression. These conclusions are unchanged when other metrics of urbanization (i.e., Human Influence Index and imperious surface cover) and indices of local adaptation (i.e., local − nonlocal_Rural_ and local − nonlocal_Urban_) are used (Figures S4-S6).

### Fitness and the Broader Microbiome

Variation in the microbiome community did not influence fitness for either white clover or rhizobia. There was no main effect of *Rhizobium* abundance or *Rhizobium* abundance × N interaction on any of the four fitness estimates (Table S3, Figure 5); using *Rhizobium* relative abundance gave similar results (Table S4, Figure S7). Local adaptation was unrelated to microbiome inoculant dissimilarity (Table S5, Figure 5) or root bacterial composition (Table S6, Figure S8).

**Figure 5:**
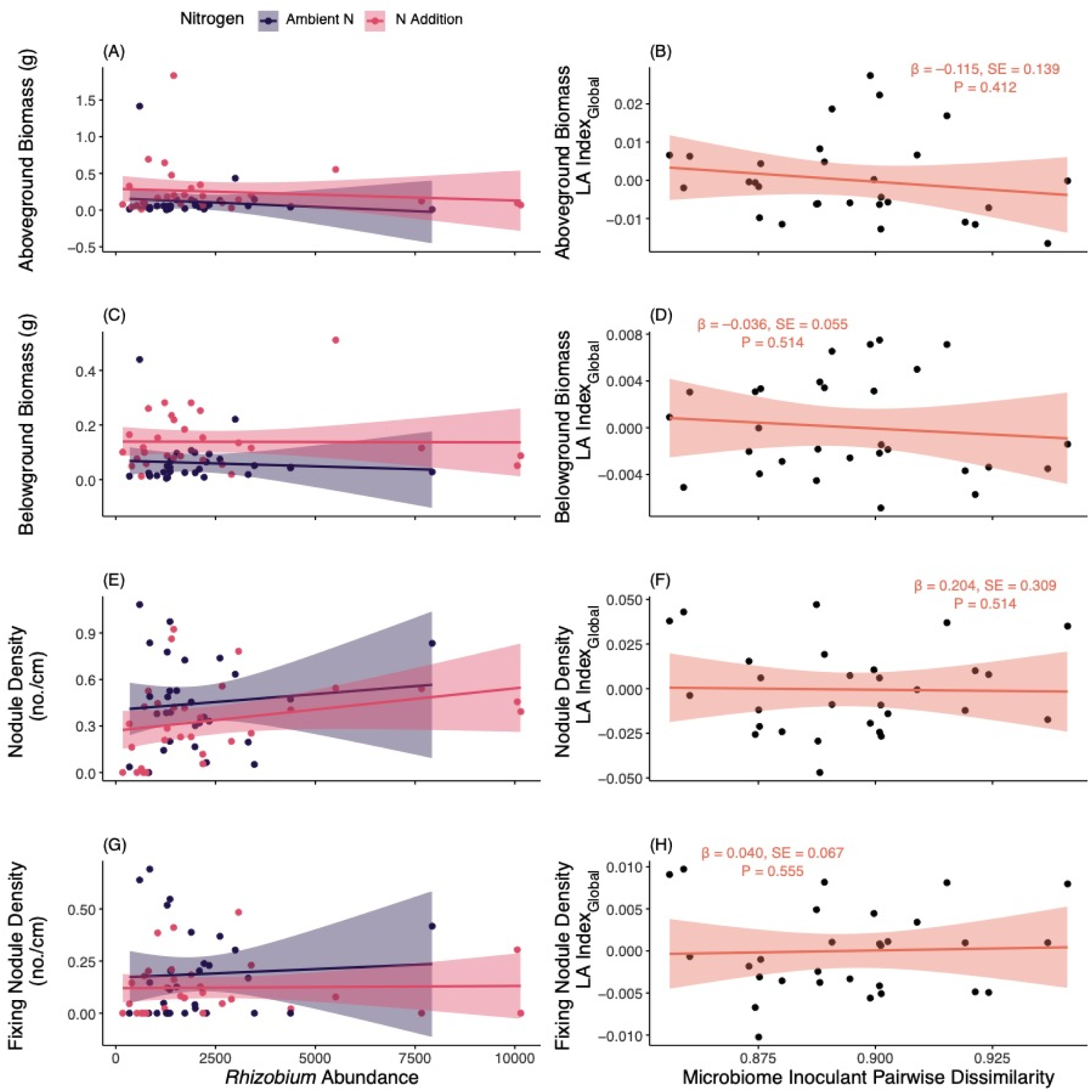
The relationship between fitness estimates and *Rhizobium* abundance (left column) and local adaptation by pairwise microbiome dissimilarity (right column) for aboveground biomass (A and B), belowground biomass (C and D), nodule density (E and F), and fixing nodule density (G and H). Lines represent lines-of-best-fit (± 95% confidence interval), with separate lines for each nitrogen treatment (ambient N = purple, N addition = pink) for the *Rhizobium* abundance models. Inset text for the left column indicates the variables of interest in the model. Influential factors and interactions were identified based on effect sizes and are in bold (η^2^_P_ range = 0.081-0.175). Inset text for the right column provides the slope parameter (β), standard error (SE), and P-value for each linear regression. Detailed test statistics are provided in Table S3 (panels A, C, E, G) and Table S5 (panels B, D, F, H).

*Rhizobium* abundance was influenced by the N treatment and an interaction between microbiome and N treatments (Table S7, Figure S9). Root bacterial community composition did not vary by microbiome inoculant, N, or the interaction (Table S6, Figure S8). Community composition was more variable among local microbiome inoculants than among either the nonlocal_Rural_ or nonlocal_Urban_ inoculants or the experimental controls (Figure S9), and *Rhizobium* abundances were similar among all microbiome inoculants (Cohen’s *d* = 0.015-0.158, Figure S10).

## Discussion

Our experiment documents a mosaic of local adaptation in the mutualism between white clover and rhizobia along an urbanization gradient, a conclusion supported by four key results. First, both white clover and rhizobia had greater mean fitness in their local microbiomes relative to the nonlocal microbiomes despite extensive variation among populations. Second, fitness responses were unrelated to *Rhizobium* abundance and local adaptation was unrelated to the broader bacterial microbiome in the soil inoculants or root endosphere. Together, these results suggest local adaptation is driven by local evolution of *Rhizobium* and not ecological changes in *Rhizobium* populations or microbiome composition. Third, N addition did not consistently mediate the effects of local adaptation in white clover or rhizobia, although N did influence the presence of reciprocal local adaptation between white clover and rhizobia, whereby the strength of local adaptation was positively correlated between white clover and rhizobia in the presence of high N. Finally, we did not find a consistent relationship between measures of urbanization and local adaptation, suggesting urban environmental change did not alter patterns of local adaptation. Below, we further discuss these results in the context of local adaptation. legume-rhizobia coevolution, and urban evolution.

### Local adaptation between white clover and rhizobia

Both white clover and rhizobia were locally adapted, although effects of local adaptation were more frequent and stronger for rhizobia. One explanation for this asymmetry in local adaptation is stronger selection on rhizobia compared to white clover. *Rhizobium* abundance and fitness is higher when cooperating with plant hosts rather than living in the soil as a saprophyte (Denison & Kiers, 2004). In contrast, white clover’s mutualism with rhizobia is facultative and dependent on the N environment, and thus selection to maintain the mutualism may be weaker (D. E. Reid et al., 2011). Another explanation is that the extensive gene flow among white clover populations within Toronto (Santangelo et al., 2022) and nearby cities (Johnson et al., 2018) could counteract the effects of local adaptation (Gomulkiewicz et al., 2007; Thompson, 2005). The population structure for rhizobia has not been characterized in this system, but a separate study found that populations of *R. leguminosarum* were genetically differentiated at spatial scales from ∼4 km to ∼350 km (Van Cauwenberghe et al., 2014), which is within the range of our urbanization gradient (∼50 km). Moreover, Van Cauwenberghe *et al*. (2016) observed genetic differentiation and associated selection mosaics contributed to local adaptation between rhizobia and their host plants. Evaluating the population structure for rhizobia is an important next step to understand the balance of selection and gene flow and its role in shaping local adaptation in our system (Gomulkiewicz et al., 2007; Hoeksema, 2010; Thompson, 2005; Wadgymar et al., 2022).

While there was an overall pattern of local adaptation, there was large variation among populations in the strength and direction of local adaptation and even maladaptation. This population × microbiome treatment interaction was driven by changes in the rank order of populations between treatments for aboveground biomass, nodule density, and fixing nodule density. Changes in rank order indicate that the relative-fitness of populations differ among environments (Wade, 2007), which can maintain genetic variation among populations when a single genotype is not the most fit in all environments (Bell et al., 2021; Christiansen, 1975; Dempster, 1955; Gomulkiewicz & Kirkpatrick, 1992). These tradeoffs could have contributed to the observed locally-adapted and locally-maladapted populations in our study.

### Microbe-mediated adaptation

By sequentially examining how the microbiome inoculants could contribute to local adaptation, we determined that *Rhizobium* evolution - not changes in *Rhizobium* abundance or microbiome community composition - likely explains local adaptation in white clover and rhizobia. Fitness responses of white clover and rhizobia were unrelated to *Rhizobium* abundance in the root endosphere. As only *R. leguminosarum* can initiate nodulation on white clover (Andrews & Andrews, 2017), our results suggest that nodulation depended on having the ‘right’ *Rhizobium* partner (i.e., genotype), which is consistent with the findings from other studies (Burghardt et al., 2022; Heath, 2010; Heath & Tiffin, 2007; Van Cauwenberghe et al., 2016). We found no relationships between either microbiome inoculant dissimilarity or root endosphere composition and local adaptation, indicating that the broader bacterial microbiome did not affect local adaptation. These findings contrast with recent studies that demonstrated adaptation to local microbiomes increases fitness (O’Brien et al., 2020; O’Brien, Yu, et al., 2022), and that dissimilarity between local and nonlocal inoculants modulates fitness responses (O’Brien, Laurich, et al., 2022). Importantly, rhizobia density in inoculants could also confound nodulation (Batstone et al., 2020; Burghardt et al., 2022), but our microbiome inoculants had similar densities of *Rhizobium* (Figure S10).

### Coevolution or correlated local adaptation?

We detected a positive correlation in the strength of local adaptation between white clover and rhizobia, which is consistent with interspecific coevolution between mutualists. While our results are consistent with a history of coadaptation resulting from coevolution, we note several qualifications with this interpretation. First, this correlation was only detected between aboveground biomass and nodule density under N addition, so whether or not a signature of coevolution is detected depends on the traits measured and the environment. Second, the presence or absence of a correlation does not necessarily implicate coevolution (Janzen, 1980; Nuismer et al., 2010). For example, spurious correlations can evolve in the absence of reciprocal biotic selection if only one of the interacting species is evolving or abiotic factors favour matching trait optima (Gomulkiewicz et al., 2007; Nuismer et al., 2010; Thompson, 2005). Alternatively, the absence of a correlation can arise despite ongoing coevolution because traits could be mismatched (Gomulkiewicz et al., 2007; Nuismer et al., 2010; Thompson, 2005). Further investigation of potential coevolution between white clover and rhizobia in urban environments requires characterizing selection acting on both white clover and rhizobia, and ideally genomic signatures of selection on genes associated with the mutualism (Batstone, Burghardt, et al., 2022; Batstone, Lindgren, et al., 2022; Epstein et al., 2023; Gomulkiewicz et al., 2007; Hoeksema, 2010).

## Conclusion

Our experiment contributes to the expanding literature on urban evolutionary ecology. Previous studies of mutualisms have predominantly focused on plant-pollinator interactions and adaptation for only one of the interacting species (Irwin *et al*. 2018; Theodorou *et al*. 2018; Rivkin *et al*. 2020; but see Brans *et al*. 2022). Moreover, studies evaluating local adaptation in urban environments have focused primarily on adaptation to abiotic factors, such as temperature (Brans et al., 2017; Martin et al., 2021) and pollutants (N. M. Reid et al., 2016; Weston et al., 2021). Anthropogenic impacts on legume-rhizobia symbioses have been evaluated in both agricultural (Lau et al., 2022; Weese et al., 2015) and more recently in urban (Forrester & Ashman, 2018; Regus et al., 2017; Wendlandt et al., 2022) environments. Nitrogen addition can lead to the evolution of less-cooperative rhizobia (Lau et al., 2022; Regus et al., 2017; Weese et al., 2015) but legumes can still maintain beneficial associations with rhizobia despite N addition and deposition (Forrester & Ashman, 2018; Wendlandt et al., 2022). Our results show that species can be locally adapted in urban environments, and that the mutualism between white clover and rhizobia can persist despite increased N (MurrayLStoker & Johnson, 2021). As plant-microbe interactions are essential for ecosystem functioning (Bardgett & van der Putten, 2014; van der Heijden et al., 2008), it is necessary to understand how microbiomes contribute to adaptation and ecosystem processes in anthropogenically-disturbed environments (Nugent & Allison, 2022).

## Acknowledgements

We thank Elana Maria, Oliva Toth, Samer El-Galmady, Wendy Van Drunen, Hind Emad, Kelly Murray-Stoker, Zain Nassrullah, and Narindra Persaud for assistance with fieldwork, Elana Maria and Alanah Joyce for help with lab work, and Joan Lee, Brenda Pitton, and the UTM Facilities staff for experiment setup. Comments from Megan Frederickson, John Stinchcombe, Sophie Breitbart, Lucas Albano, and Aude Caizergues greatly improved the manuscript. This work was funded by an NSERC Discovery Grant, Canada Research Chair, and E.W.R. Steacie Fellowship to M. T. J. Johnson and an American Society of Naturalists Student Research Award to D. Murray-Stoker.

## Author Contributions

DMS and MTJJ designed the experiment. DMS collected and analyzed the data, with input from MTJJ. MTJJ and DMS secured the funding. DMS led the writing of the manuscript, with support from MTJJ.

## Data Availability Statement

Data and code are available on Zenodo (<u>10.5281/zenodo.8253951</u>). Sequences are available at NCBI under the BioProject accession <u>PRJNA1005557</u>.

