## Appendix S1 for "Mosaic of local adaptation between white clover and rhizobia along an urbanization gradient"

**Supplemental Methods**

*Generation of White Clover F_1_ Seeds*

We grew field-collected seeds in the greenhouse to produce outcrossed F_1_ populations to minimize parental effects. The 30 focal white clover populations for this experiment were a subset from those sampled by Murray-Stoker and Johnson (2021). At each population, we collected ~10 ripe infructescences, with plants separated by a minimum of 5 m to avoid collecting the same clone.

Seeds were returned to the University of Toronto Mississauga, scarified with sandpaper, and germinated in the dark on dampened filter paper for 48 h at 20°C. Germinated seeds were then planted in 100 mL pots filled with Pro-Mix LP15 potting soil. Plants were grown for 12 weeks, with weeks 1-7 set to an 18:6 light:dark cycle with 350 µmol of light, day temperature at 25°C, night temperature at 14°C, and humidity at 50%. To facilitate fruit and seed development, weeks 8-12 were maintained at an 18:6 light:dark cycle, but light was maximized to 700 µmol, day temperature at 21°C, night temperature at 13°C; humidity was maintained at 50%. Each plant received 3-5 Nutricote Total 14-13-13 (N-P-K) pellets after the first true leaf expanded. Seeds were generated from a random panmictic cross among 5-10 parent plants from each source population. Pollen was transferred between plants by hand using a paintbrush. Seeds were collected from mature fruits after week 12 and stored in a refrigerator at 4°C.

*Sample Preparation for DNA Extraction*

*Roots:* We selected 60 root samples for sequencing (10 populations × 3 microbiome treatments [local, nonlocal_Rural_, nonlocal_Urban_] × 2 N treatments [ambient N and N addition] = 60 plants). All root mass was collected from each plant, dried (at least 72 h at 60°C) to measure belowground biomass, and placed in 15 mL Falcon tubes for washing in sterile water using a Vortex at maximum speed for 10 s. We also sonicated root samples for 10 m at 60 Hz to facilitate physical removal of any remaining soil. Roots were freeze-dried (Freeze Dryer Epsilon 2-6D LSCplus) and then ground and homogenized (QIAGEN TissueLyser II). We weighed approximately 80 mg of roots for each sample (mean ± SE = 80.70 mg ± 8.91 mg), with samples stored in the freezer at −80°C until DNA extraction.

*Soil:* All 40 microbiome inoculants (30 focal, 5 nonlocal_Rural_, 5 nonlocal_Urban_) plus the 6 experimental controls (3 ambient N, 3 N addition) were prepared for sequencing. We weighed 250 mg of soil and 80 mg of roots, with root samples washed, freeze-dried (Freeze Dryer Epsilon 2-6D LSCplus), and homogenized (QIAGEN TissueLyser II) to facilitate DNA extraction. Soil samples were removed from a −80°C freezer and thawed for ~12 hours in a refrigerator at 4°C, and approximately 250 mg (wet weight) of each soil sample was weighed for DNA extraction.

*DNA Extraction & Sequencing*

We extracted DNA from each soil and root sample using the QIAGEN DNeasy PowerSoil Pro Kit following the manufacturer’s protocol. We targeted the V4 hypervariable region of 16S for sequencing(primers 515f and 806r; Walters *et al.* 2016), with peptide nucleic acid (PNA) clamps to inhibit amplification of plant plastid and mitochondrial DNA (Lundberg *et al.* 2013; Fitzpatrick *et al.* 2018). Soil DNA was normalized to approximately 50 ng/µL (50.027 ng/µL ± 2.987 ng/µL), but root DNA was not normalized due to low and variable concentrations of extracted DNA (28.072 ng/µL ± 13.959 ng/µL) that could have been confounded by both microbial and plant DNA; a Nanodrop spectrophotometer was used to quantify DNA concentrations. All samples were amplified and sequenced by the Surette Lab at McMaster University on an Illumina MiSeq with 2 x 300 paired-end reads.

*Statistical Analyses*

*Model selection:* We employed a model fitting and information theoretic-based selection procedure to optimize random effect structures for analysis. We started with a base model (Model 1; see list of model structures below), with alternative models (Models 2 and 3) only selected if there was a clear improvement in model performance (i.e., ΔAIC_C_ > 2, model weights > 0.70). Model 1 was our base model with random intercepts, and it was fitted as:

Response = Microbiome + Nitrogen + Microbiome:Nitrogen

+ (1 | Microbiome:Population)

+ (1 | Nitrogen:Population)

+ (1 | Microbiome:Nitrogen:Population)

+ (1 | Population)

+ (1 | Block)

Model 2 added random intercepts for the microbiome and nitrogen treatments, and was fitted as:

Response = Microbiome + Nitrogen + Microbiome:Nitrogen

+ (Microbiome | Population)

+ (Nitrogen | Population)

+ (Microbiome:Nitrogen |Population)

+ (1 | Population)

+ (1 | Block)

Model 3 was the simplified model with only random intercepts for population and block, fitted as:

Response = Microbiome + Nitrogen + Microbiome:Nitrogen

+ (1 | Population)

+ (1 | Block)

If Model 1 was identified as the best-fitting model after the first round of model selection, we then systematically fitted and compared subsets of Model 1 to determine if dropping specific random effect terms improved model fit (see below for detailed procedure). When dropping interaction terms within the random effects, we ensured lower-order components were retained in the model structure.

Model 1 was our base model and allowed us to test for the main effects of microbiome and nitrogen on fitness estimates in addition to the two- and three-way interactions between microbiome and nitrogen with population identity. Each of these three variables (i.e., microbiome treatment, nitrogen treatment, and population) and their interactions were relevant to the research questions for the experiment. Each of the Model 1 subsets dropped a specific interaction: Model 1.1 dropped the three-way interaction between microbiome treatment, nitrogen treatment, and population [i.e., (1 | Microbiome:Nitrogen:Population) was dropped from the model]; Model 1.2 dropped the three-way interaction between microbiome treatment, nitrogen treatment, and population plus the component two-way interaction between microbiome treatment and population [i.e., (1 | Microbiome:Population) and (1 | Microbiome:Nitrogen:Population) were dropped from the model]; and Model 1.3 dropped the three-way interaction between microbiome treatment, nitrogen treatment, and population plus the component two-way interaction between nitrogen treatment and population [i.e., (1 | Nitrogen:Population) and (1 | Microbiome:Nitrogen:Population) were dropped from the model]. Each of these model subsets still allowed us to test variables of interest while identifying model structure that could optimize statistical power and model performance.

In our comparison of Model 1 and its subsets, we selected the model with the lowest AIC_C_ as the optimal model, even if other models were ‘statistically equivalent’ (i.e., ΔAIC_C_ < 2). While this is in contrast to common statistical practice (Johnson & Omland 2004; Grueber *et al.* 2011), our objective with this model selection procedure was to optimize random effect structures while retaining models that tested the variables of interest.

*Microbiome × Nitrogen Analyses:* We analyzed variation in fitness responses for white clover (aboveground biomass, belowground biomass) and rhizobia (nodule density, fixing nodule density) by microbiome treatment (local, nonlocal_Rural_, and nonlocal_Urban_), nitrogen treatment (ambient N and N addition), and the two-way interaction using linear mixed-effects models with the `lme4` (Bates *et al.* 2015)and `lmerTest` (Kuznetsova *et al.* 2017) packages. We used the model structures identified through our model selection procedure described above. Based on our model selection process, aboveground biomass was evaluated using Model 1.1 (ΔAIC_C_ = 0.934, model weight = 0.420; interpretation of results for Model 1 and Model 1.1 were similar), belowground biomass with Model 3 (ΔAIC_C_ = 5.399, model weight = 0.937), nodule density with Model 1 (ΔAIC_C_ = 4.318, model weight = 0.864), and fixing nodule density with Model 1.1 (ΔAIC_C_ = 1.627, model weight = 0.577).

Influence of fixed effects was estimated using ANOVAs with Type III sums-of-squares, with degrees of freedom estimated using the Kenward-Roger method (Kenward & Roger 1997). Effect sizes for the fixed effects were calculated as partial eta-squared (η^2^_P_), which explains the amount of variation explained by the model term (Cohen 1973). Influence of random effects was estimated using likelihood ratio tests, with the effect size for the random effect terms calculated as the intra-class correlation coefficient (ICC; Gelman & Hill 2006; Nakagawa *et al.* 2017), which measures the amount of variation explained by random effect term. Post-hoc comparisons were conducted using estimated marginal means weighted by cell frequencies, and effect sizes for contrasts were calculated as Cohen’s *d* using the `eff_size()` function; estimated marginal means and contrasts were calculated using the `emmeans` package (Lenth *et al.* 2022). Both η^2^_P_ and ICC can be interpreted similarly to partial R^2^ (Cohen 1973; Gelman & Hill 2006; Nakagawa *et al.* 2017).

*Local Adaptation Indices:* Local adaptation indices were quantified to evaluate the degree of local adaptation. Three separate local adaptation indices were calculated: (1) local − nonlocal_Global_, (2) local − nonlocal_Rural_, and (3) local − nonlocal_Urban_, wherein the estimate of the respective nonlocal treatment [global (nonlocal_Rural_ and nonlocal_Urban_), nonlocal_Rural_, and nonlocal_Urban_) was subtracted from the estimate of the local treatment for each of the four fitness response variables. Positive values indicated greater fitness in the local treatment while negative values indicated greater fitness in the nonlocal treatment. Local adaptation indices were calculated by refitting all linear mixed-effects models for the microbiome × nitrogen analyses by using the full model structure:

Response ~ Microbiome + Nitrogen + Microbiome:Nitrogen

+ (1 | Microbiome:Population)

+ (1 | Nitrogen:Population)

+ (1 | Microbiome:Nitrogen:Population)

+ (1 | Population)

+ (1 | Block)

We then calculated the best linear unbiased predictors (BLUPs) for the (1 | Microbiome:Nitrogen:Population) term to quantify the local adaptation indices, with BLUPs calculated for each population, microbiome, and nitrogen treatment combination.

*Correlations between Local Adaptation Indices:* We correlated local adaptation indices to test for potential evidence of coevolution. For each local adaptation index, we made 6 comparisons: (1) aboveground biomass and belowground biomass, (2) nodule density and fixing nodule density, (3) aboveground biomass and nodule density, (4) aboveground biomass and fixing nodule density, (5) belowground biomass and nodule density, (6) belowground biomass and fixing nodule density. Comparisons allowed us to test for correlations within (e.g., aboveground and belowground biomass, nodule density and fixing nodule density) and between (e.g., aboveground biomass and nodule density) partners in the mutualism. For each comparison, we also compared the Pearson correlation coefficient between N treatments to examine if N addition mediated the relationship. In total, there were 12 comparisons made for each local adaptation index (6 comparisons × 2 N treatments = 12). It is important to note that the presence or absence of a correlation is not definitive evidence of coevolution (Gomulkiewicz *et al.* 2007; Nuismer *et al.* 2010).

*Urbanization & Local Adaptation:* To test how urbanization affects the strength of local adaptation, we regressed local adaptation indices against measures of urbanization. We first calculated three measures of urbanization: distance from the city center, mean impervious surface cover (ISC), and the human influence index (HII). Distance from the urban center was calculated for each site as the distance on an ellipsoid (i.e., the geodesic distance) using the `distm()` function in the geosphere package (Hijmans 2019). Coordinates for the urban center were selected as the Toronto City Hall (43.651536, -79.383276). Impervious surface cover (ISC) was estimated using the 30-m resolution Global Man-Made Impervious Surface raster dataset (Brown de Colstoun *et al.* 2017), with ISC values averaged within a 250-m buffer surrounding each population. Human influence index (HII) is a globally-distributed dataset (1-km grid cells) containing data on human population density, land use, infrastructure, and human access in a composite variable of HII (Wildlife Conservation Society - WCS & Center for International Earth Science Information Network - CIESIN - Columbia University 2005). We calculated ISC and HII values for each population using functions in the `raster` package (Hijmans *et al.* 2023).

After calculating measures of urbanization, we tested for effects of urbanization on local adaptation using linear regressions. We fitted the linear regressions as:

Local Adaptation Index = Intercept + x_i_ + *e*_i_

where a local adaptation index (local − nonlocal_Global_, local − nonlocal_Rural_, and local − nonlocal_Urban_) was the response, x_i_ was the urbanization metric (distance from the city center, mean ISC, or HII), and *e*_i_ was the residual error. A full model with all predictor variables was not fitted because multicollinearity between predictor variables (variance inflation factors > 5–10) made it difficult to disentangle the effects of individual variables and would have violated model assumptions. We standardized (i.e., mean = 0, standard deviation = 1) all urbanization metric predictor variables prior to analysis to allow for direct comparisons across models. All model assumptions were evaluated graphically, and we evaluated the slope parameter, standard error of the slope estimate, P-value, and variance explained by the model (R^2^) to determine the evidence for an effect of urbanization on the strength of local adaptation.

*Microbiome Analyses:* We processed demultiplexed reads using the DADA2 pipeline (Callahan *et al.* 2016) to prepare the samples for further analyses. As the root and soil sequences were generated from different sequencing runs, we put the sequences though separate iterations of the DADA2 pipeline to have a more accurate error rate. We first examined quality profiles of the forward and reverse reads, and we trimmed reads when the Phred quality (Q) score dropped (i.e., Q < 30). For the root sequences, forward reads were trimmed at 225 bp and reverse reads were trimmed at 215 bp. For the soil sequences, forward reads were trimmed at 250 bp and reverse reads were again trimmed at 215 bp. All sequences had the first 10 bp removed from the sequence due to low quality scores. We also filtered any sequences with ambiguous nucleotide assignments, with any instance of a Q-score < 2, and/or sequences with more than 2 expected errors (Callahan *et al.* 2016).

After sequences were trimmed and filtered, we used the DADA2 algorithm to learn the error rates (trained on 6 samples or 10% of the samples) and then inferred bacterial taxa using DADA2 sample inference, which infers taxa as amplicon sequence variants (ASVs). DADA2 uses the sequences and error rates supplied by the user to identify probable sequences instead of PCR or sequencing artifacts. As a result, DADA2 has very high accuracy for identifying taxa (but see critiques by Nearing *et al.* 2018). For the roots, there were 471 sequence variants inferred from 23172 input unique sequences for the forward reads and 691 sequence variants inferred from 15971 input unique sequences for the reverse reads. For the soil, there were 608 sequence variants inferred from 52092 input unique sequences from the forward reads and 1379 sequence variants from 32376 input unique sequences for the reverse reads. Once ASVs were identified, we merged the forward and reverse reads and removed chimeras. We then assigned taxonomy to individual ASVs using the RDP naïve Bayesian classifier (Wang *et al.* 2007) and the Ribosomal Database Project training set 18 (Cole *et al.* 2014) implemented in DADA2.

First, we compared fitness estimates by rhizobia abundance. Linear models were fitted as:

y_i_ = Intercept + Rhizobia Abundance + Nitrogen + Rhizobia Abundance:Nitrogen

+ Sample Reads + *e*_i_

where for each fitness estimate y_i_ (aboveground biomass, belowground biomass, nodule density, or fixing nodule density), we evaluated the main effects of rhizobia abundance, nitrogen treatment (ambient N and N addition), and the two-way interaction; *e*_i_ was the residual error. We also fitted the number of sample reads to control for uneven sequencing depth across samples. Moreover, we fitted an identical set of models, but we replaced rhizobia abundance with rhizobia relative abundance. Assumptions for each linear model were inspected graphically, and influence of fixed effects was evaluated using Type III sums-of-squares.

Second, we compared rhizobia abundance and relative abundance by microbiome and nitrogen treatments using ANOVAs. Separate models were fitted for rhizobia abundance and rhizobia relative abundance, with each ANOVA fitted with the form:

Abundance Estimate = Intercept + Microbiome_Global_ + Nitrogen

+ Microbiome_Global_:Nitrogen + *e*_i_

where either rhizobia abundance or rhizobia relative abundance was the abundance estimate, and we evaluated the main effects of microbiome_Global_ (local and nonlocal_Global_), nitrogen treatment (ambient N and N addition), and the two-way interaction; *e*_i_ was the residual error. Rhizobia abundance was fitted using a generalized linear model with a Poisson distribution, while rhizobia relative abundance was fitted using a linear model. All model assumptions were inspected graphically. Influence of experimental treatments was estimated using ANOVAs with Type III sums-of-squares, with Wald 𝜒^2^ tests for rhizobia abundance and F-tests for rhizobia relative abundance. Similarly, effect sizes for predictors in the rhizobia abundance model were quantified as Cohen’s *w* (Cohen 1988), while effect sizes for predictors in the rhizobia relative abundance model were calculated as η^2^_P_. Higher values for Cohen’s *w* indicate a stronger association between the predictor variable and response (Cohen 1988).

Third, we related the broader root bacterial microbiome composition to fitness estimates. We calculated a square root-transformed Bray-Curtis dissimilarity matrix among root communities. We then conducted a PERMANOVA with 10,000 permutations on this distance matrix to test for differences in community composition by microbiome and nitrogen treatments using `adonis2()` (Oksanen *et al.* 2020), with the model fitted as:

Distance Matrix = Microbiome_Global_ + Nitrogen + Microbiome_Global_:Nitrogen

where we evaluated the main effects of microbiome_Global_ (local and nonlocal_Global_), nitrogen treatment (ambient N and N addition), and the two-way interaction. We also performed a non-metric multidimensional scaling ordination using `monoMDS()` (Oksanen *et al.* 2020). Relationships between community composition and fitness estimates were determined using the `envfit()` function with 1000 permutations (Oksanen *et al.* 2020), and this analysis shows the direction and strength of the relationship (measured as R^2^) between the fitness estimates and community composition on the ordination.

Finally, we evaluated community composition between all microbiome inoculants using a principal coordinates analysis (PCoA). Each of the microbiome inoculants (local, nonlocal_Rural_, nonlocal_Urban_) was included in the PCoA, including the 6 control pots (3 ambient N, 3 N addition), to evaluate soil bacterial community composition. After the PCoA was performed, we visualized separation of communities by community type (i.e., soil inoculant, ambient N control, N addition control) in ordination space. We conducted the PCoA on Euclidean distances using the `capscale()` function (Oksanen *et al.* 2020). In addition to comparisons of bacterial community composition, we compared the abundance of rhizobia by community type using a generalized linear model with a Poisson distribution. The model was fitted as:

*Rhizobium* Abundance ~ Intercept + Community Type + *e*

where *Rhizobium* abundance (number of reads identified to *Rhizobium*) was the response, community type was the predictor, and *e* was the residual error. Model assumptions were inspected graphically, and influence of community type was estimated using an ANOVA with Type III sums-of-squares and Wald 𝜒^2^ tests. The effect size for community type was quantified as Cohen’s *w*.

*Statistical Software:* All of the above analyses were performed using R (version 4.2.2; R Core Team 2022) in the RStudio environment (version 2022.07.2; RStudio Team 2022). Graphical evaluation of model fits was conducted using the `performance` package (Lüdecke *et al.* 2021). Effect sizes were calculated using the `effectsize` (η^2^_P_ and Cohen’s *w*; Ben-Shachar *et al.* 2020), `performance`(ICC; Lüdecke *et al.* 2021), and `emmeans` (Cohen's *d*; Lenth *et al.* 2022) packages. Microbiome analyses were facilitated by the `phyloseq` (McMurdie & Holmes 2013) and `tidyamplicons` packages (Wittouck 2023). Data management and figure creation were facilitated using the `tidyverse` (Wickham *et al.* 2019) and `ggpubr` (Kassambara 2020) packages. We did not use arbitrary thresholds of statistical significance to categorize evidence. Instead, we followed recommendations for statistical best practices by focusing on estimates, variation of the estimates, and effect sizes to identify the strength of evidence (Carver 1978; Nakagawa & Cuthill 2007; Wasserstein & Lazar 2016; Gelman & Hennig 2017; McShane *et al.* 2019; Wasserstein *et al.* 2019; Berner & Amrhein 2022).

**Supplemental Tables**

**Table S1:** Contrasts from the linear mixed-effects models testing the effects of microbiome (local, nonlocal_Rural_, nonlocal_Urban_), nitrogen (ambient N and N addition), and the two-way interaction on measures of plant and rhizobia fitness. We calculated contrasts using estimated marginal means, with effect sizes for the contrasts measured as Cohen’s *d* and the associated 95% confidence interval. We report the inequality for each contrast alongside the effect size (Cohen’s *d*), with confidence intervals are reported in square brackets.

| **Fitness Estimate** | **Microbiome Contrast** | | |  | **Nitrogen Contrast** |
| --- | --- | --- | --- | --- | --- |
|  | **Local vs. Nonlocal_Rural_** | **Local vs. Nonlocal_Urban_** | **Nonlocal_Urban_ vs. Nonlocal_Rural_** |  | **N Addition vs Ambient N** |
| Aboveground Biomass | Local > Nonlocal_Rural_  *d* = 0.136 [−0.073, 0.345] | Local > Nonlocal_Urban_  *d* = 0.253 [0.044, 0.463] | Nonlocal_Urban_ < Nonlocal_Rural_  *d* = −0.117 [−0.326. 0.091] |  | N Addition > Ambient N  *d* = 1.222 [1.029, 1.415] |
| Belowground Biomass | Local ≈ Nonlocal_Rural_  *d* = 0.012 [−0.155, 0.179] | Local > Nonlocal_Urban_  *d* = 0.172 [0.005, 0.339] | Nonlocal_Urban_ < Nonlocal_Rural_  *d* = −0.160 [−0.327, 0.006] |  | N Addition > Ambient N  *d* = 1.356 [1.205, 1.506] |
| Nodule Density | Local > Nonlocal_Rural_  *d* = 0.549 [0.289, 0.809] | Local > Nonlocal_Urban_  *d* = 0.332 [0.073, 0.591] | Nonlocal_Urban_ > Nonlocal_Rural_  *d* = 0.217 [−0.042, 0.475] |  | N Addition < Ambient N  *d* = −0.331 [−0.520, −0.143] |
| Fixing Nodule Density | Local > Nonlocal_Rural_  *d* = 0.482 [0.257, 0.707] | Local > Nonlocal_Urban_  *d* = 0.352 [0.128. 0.575] | Nonlocal_Urban_ > Nonlocal_Rural_  *d* = 0.130 [−0.092. 0.353] |  | N Addition > Ambient N  *d* = −0.215 [−0.404, −0.025] |

**Table S2:** Summary of the linear mixed-effects models testing the effects of microbiome (local and nonlocal_Global_), nitrogen (ambient N and N addition), and the two-way interaction on measure of plant and rhizobia fitness. We provide the numerator degrees of freedom (NumDF), denominator degrees of freedom (approximated using the Kenward-Roger method (DenDF), F-statistics calculated from type III sums-of-squares, P-values, and the effect size (partial eta-squared, η^2^_P_) for each fixed effect term in the ANOVAs. We also provide the χ^2^ statistic, P-values from likelihood ratio tests, and the effect size (intra-class correlation coefficient, ICC) for each random effect term, with random effect term and statistics in italics. The variance explained by the fixed effects (R^2^_Marginal_) and combined fixed and random effects (R^2^_Conditional_) is provided for each model.

| **Term** | **NumDF** | **DenDF** | **F/**χ^2^ | **P-value** | **η^2^_P_/ICC** |
| --- | --- | --- | --- | --- | --- |
| **Aboveground Biomass**  **(R^2^_Marginal_ = 0.230, R^2^_Conditional_ = 0.394)** | | | | | |
| Microbiome | 1 | 28.468 | 4.057 | 0.054 | 0.125 |
| Nitrogen | 1 | 32.642 | 173.887 | < 0.001 | 0.842 |
| Microbiome × Nitrogen | 1 | 766.967 | 0.545 | 0.461 | 0.001 |
| *Microbiome × Population* |  |  | 5.416 | 0.020 | 0.044 |
| *Nitrogen × Population* |  |  | 4.118 | 0.042 | 0.037 |
| *Population* |  |  | 7.867 | 0.005 | 0.120 |
| *Block* |  |  | 4.182 | 0.041 | 0.012 |
| **Belowground Biomass**  **(R^2^_Marginal_ = 0.288 R^2^_Conditional_ = 0.376)** | | | | | |
| Microbiome | 1 | 803.909 | 1.520 | 0.218 | 0.002 |
| Nitrogen | 1 | 805.166 | 329.021 | < 0.001 | 0.290 |
| Microbiome × Nitrogen | 1 | 806.873 | 0.255 | 0.614 | < 0.001 |
| *Population* |  |  | 46.467 | < 0.001 | 0.105 |
| *Block* |  |  | 4.654 | 0.031 | 0.014 |
| **Nodule Density**  **(R^2^_Marginal_ = 0.055, R^2^_Conditional_ = 0.263)** | | | | | |
| Microbiome | 1 | 31.390 | 18.108 | < 0.001 | 0.366 |
| Nitrogen | 1 | 35.652 | 8.400 | 0.006 | 0.191 |
| Microbiome × Nitrogen | 1 | 89.952 | 1.111 | 0.295 | 0.012 |
| *Microbiome × Population* |  |  | < 0.001 | ≈ 1.000 | < 0.001 |
| *Nitrogen × Population* |  |  | < 0.001 | ≈ 1.000 | < 0.001 |
| *Microbiome × Nitrogen x Population* |  |  | 26.256 | < 0.001 | 0.158 |
| *Population* |  |  | 4.791 | 0.029 | 0.058 |
| *Block* |  |  | 0.753 | 0.385 | 0.004 |
| **Fixing Nodule Density**  **(R^2^_Marginal_ = 0.041, R^2^_Conditional_ = 0.162)** | | | | | |
| Microbiome | 1 | 29.102 | 25.754 | < 0.001 | 0.469 |
| Nitrogen | 1 | 32.602 | 6.524 | 0.016 | 0.167 |
| Microbiome × Nitrogen | 1 | 769.770 | 0.672 | 0.413 | 0.001 |
| *Microbiome × Population* |  |  | 0.403 | 0.526 | 0.012 |
| *Nitrogen × Population* |  |  | 4.463 | 0.035 | 0.042 |
| *Population* |  |  | 1.834 | 0.176 | 0.039 |
| *Block* |  |  | 15.030 | < 0.001 | 0.032 |

**Table S3:** Summary of the ANCOVAs comparing fitness estimates by *Rhizobium* abundance, nitrogen (ambient N and N addition), and the two-way interaction; the number of reads was included in the models as a covariate. We provide the F-statistics calculated from Type III sums-of-squares, P-values, and effect size (partial eta-squared, η^2^_P_) for each term in the models.

| **Term** | **F** | **P-value** | **η^2^_P_** |
| --- | --- | --- | --- |
| **Aboveground Biomass** | | | |
| Abundance | 0.528 | 0.470 | 0.010 |
| Nitrogen | 4.790 | 0.033 | 0.081 |
| Abundance × Nitrogen | 0.141 | 0.709 | 0.003 |
| Number of Reads | 2.120 | 0.151 | 0.038 |
| **Belowground Biomass** | | | |
| Abundance | 0.009 | 0.924 | < 0.001 |
| Nitrogen | 11.423 | 0.001 | 0.175 |
| Abundance × Nitrogen | 0.140 | 0.709 | 0.003 |
| Number of Reads | 5.864 | 0.019 | 0.098 |
| **Nodule Density** | | | |
| Abundance | 0.741 | 0.393 | 0.014 |
| Nitrogen | 1.488 | 0.228 | 0.027 |
| Abundance × Nitrogen | 0.012 | 0.913 | < 0.001 |
| Number of Reads | 1.197 | 0.279 | 0.022 |
| **Fixing Nodule Density** | | | |
| Abundance | 0.644 | 0.426 | 0.012 |
| Nitrogen | 0.195 | 0.661 | 0.004 |
| Abundance × Nitrogen | 0.343 | 0.560 | 0.006 |
| Number of Reads | 2.072 | 0.156 | 0.037 |

Note: There was 1 numerator df and 54 residual df for each model term.

**Table S4:** Summary of the ANCOVAs comparing fitness estimates by *Rhizobium* relative abundance, nitrogen (ambient N and N addition), and the two-way interaction; the number of reads was included in the models as a covariate. We provide the F-statistics calculated from Type III sums-of-squares, P-values, and effect size (partial eta-squared, η^2^_P_) for each term in the models.

| **Term** | **F** | **P-value** | **η^2^_P_** |
| --- | --- | --- | --- |
| **Aboveground Biomass** | | | |
| Relative Abundance | 0.460 | 0.501 | 0.008 |
| Nitrogen | 3.389 | 0.071 | 0.059 |
| Relative Abundance × Nitrogen | 0.321 | 0.574 | 0.006 |
| Number of Reads | 1.292 | 0.261 | 0.023 |
| **Belowground Biomass** | | | |
| Relative Abundance | 0.271 | 0.605 | 0.005 |
| Nitrogen | 12.572 | 0.001 | 0.189 |
| Relative Abundance × Nitrogen | 0.598 | 0.443 | 0.011 |
| Number of Reads | 5.823 | 0.019 | 0.097 |
| **Nodule Density** | | | |
| Relative Abundance | 3.388 | 0.071 | 0.059 |
| Nitrogen | 0.016 | 0.901 | < 0.001 |
| Relative Abundance × Nitrogen | 1.449 | 0.234 | 0.026 |
| Number of Reads | 0.089 | 0.767 | 0.002 |
| **Fixing Nodule Density** | | | |
| Relative Abundance | 2.447 | 0.124 | 0.043 |
| Nitrogen | 0.022 | 0.918 | < 0.001 |
| Relative Abundance × Nitrogen | 1.563 | 0.217 | 0.028 |
| Number of Reads | 1.037 | 0.313 | 0.019 |

Note: There was 1 numerator df and 54 residual df for each model term.

**Table S5:** Summary of the linear models comparing the strength of local adaptation (local – nonlocal_Global_) by the pairwise Bray-Curtis dissimilarity of the local and nonlocal_Global_ microbiomes. We provide the slope ($\beta$), standard error of the slope (SE), t-statistic, P-value, and standardized slope (standardized $\beta$) for each term in the models. We also provide the adjusted R^2^ (R^2^_Adjusted_) for each model.

| **Term** | $\boldsymbol{\beta}$ | **SE** | **t** | **P-value** | **Standardized** $\boldsymbol{\beta}$ |
| --- | --- | --- | --- | --- | --- |
| **Aboveground Biomass**  **(R^2^_Adjusted_ < 0.001)** | | | | | |
| Pairwise Microbiome Dissimilarity | –0.115 | 0.139 | –0.833 | 0.412 | –0.232 |
| Number of Reads | < 0.001 | < 0.001 | –0.311 | 0.758 | –0.087 |
| **Belowground Biomass**  **(R^2^_Adjusted_ < 0.001)** | | | | | |
| Pairwise Microbiome Dissimilarity | –0.036 | 0.055 | –0.662 | 0.514 | –0.186 |
| Number of Reads | < 0.001 | < 0.001 | –0.401 | 0.692 | –0.113 |
| **Nodule Density**  **(R^2^_Adjusted_ < 0.001)** | | | | | |
| Pairwise Microbiome Dissimilarity | 0.204 | 0.309 | 0.662 | 0.514 | 0.184 |
| Number of Reads | < 0.001 | < 0.001 | 1.032 | 0.312 | 0.287 |
| **Fixing Nodule Density**  **(R^2^_Adjusted_ < 0.001)** | | | | | |
| Pairwise Microbiome Dissimilarity | 0.040 | 0.067 | 0.598 | 0.555 | 0.168 |
| Number of Reads | < 0.001 | < 0.001 | 0.638 | 0.529 | 0.179 |

Note: There was 1 numerator df and 26 residual df for each model term.

**Table S6:** Summary of the ANOVAs comparing either *Rhizobium* abundance or *Rhizobium* relative abundance by microbiome (local vs. nonlocal_Global_), nitrogen (ambient N and N addition), and the two-way interaction. We report the test statistic (χ^2^ = *Rhizobium* abundance, F = *Rhizobium* relative abundance) calculated from Type III sums-of-squares, P-value, and effect size (Cohen’s w = *Rhizobium* abundance, partial eta-squared (η^2^_P_) = *Rhizobium* relative abundance) for each term in the models.

| **Term** | **χ^2^/F** | **P-value** | **Cohen’s *w*/ η^2^_P_** |
| --- | --- | --- | --- |
| ***Rhizobium* Abundance** | | | |
| Microbiome | 0.769 | 0.380 | 0.113 |
| Nitrogen | 39.060 | < 0.001 | 0.807 |
| Microbiome × Nitrogen | 373.176 | < 0.001 | 2.494 |
| ***Rhizobium* Relative Abundance** | | | |
| Microbiome | 0.034 | 0.854 | 0.001 |
| Nitrogen | 1.097 | 0.299 | 0.020 |
| Microbiome × Nitrogen | 0.367 | 0.547 | 0.007 |

Note: There was 1 numerator df and 55 residual df for each model term.

**Table S7:** Summary of the PERMANOVA comparing root bacterial community composition by microbiome (local vs. nonlocal_Global_), nitrogen (ambient N, N addition), and the two-way interaction. We provide the F-statistic, P-value, and R^2^ for each model term. We also provide the P-value and R^2^ for vectors of the local adaptation index for each fitness estimate in relation to community composition, with these statistics in italics. We calculated P-values for the local adaptation index vectors from 1000 permutations using the `envfit` function in the `vegan` package (Oksanen et al. 2020).

| **Term** | **F** | **P-value** | **R^2^** |
| --- | --- | --- | --- |
| Microbiome | 0.856 | 0.815 | 0.015 |
| Nitrogen | 1.011 | 0.414 | 0.017 |
| Microbiome × Nitrogen | 0.719 | 0.991 | 0.012 |
| *Aboveground Biomass* |  | *0.918* | *0.004* |
| *Belowground Biomass* |  | *0.338* | *0.036* |
| *Nodule Density* |  | *0.587* | *0.018* |
| *Fixing Nodule Density* |  | *0.724* | *0.011* |

Note: There was 1 numerator df and 56 residual df for each model term the PERMANOVA.

**Supplemental** **Figures**

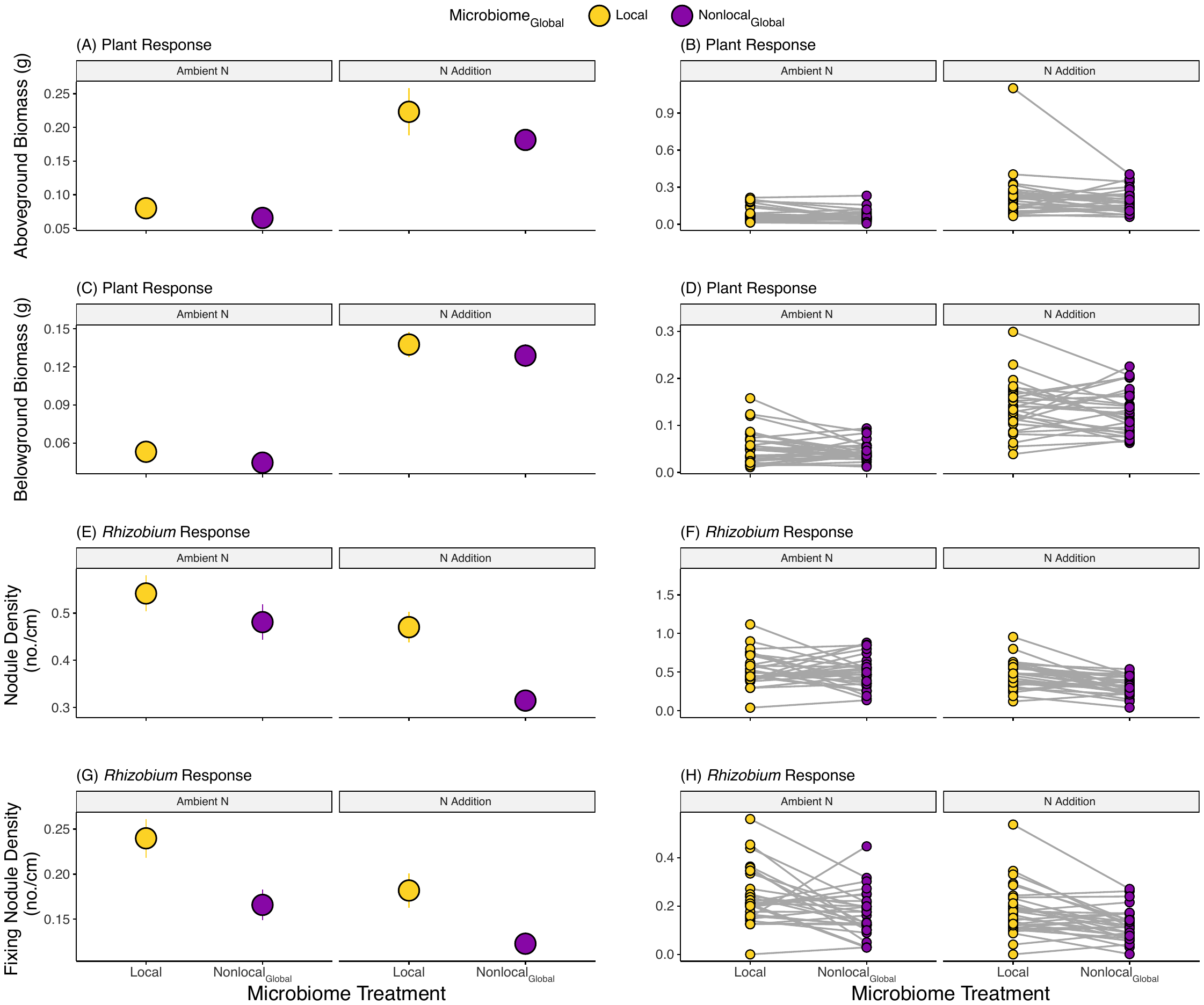

**Figure S1:** Treatment means ± 1 SE (left column) and reaction norms of population means (right column) by microbiome and nitrogen treatments for aboveground biomass (A-B), belowground biomass (C-D), nodule density (E-F), and fixing nodule density (G-H). Microbiome treatments are indicated as local (blue) and nonlocal_Global_ (purple); individual plots are faceted by nitrogen treatment (ambient N and N addition). Detailed test statistics are provided in Table S2.

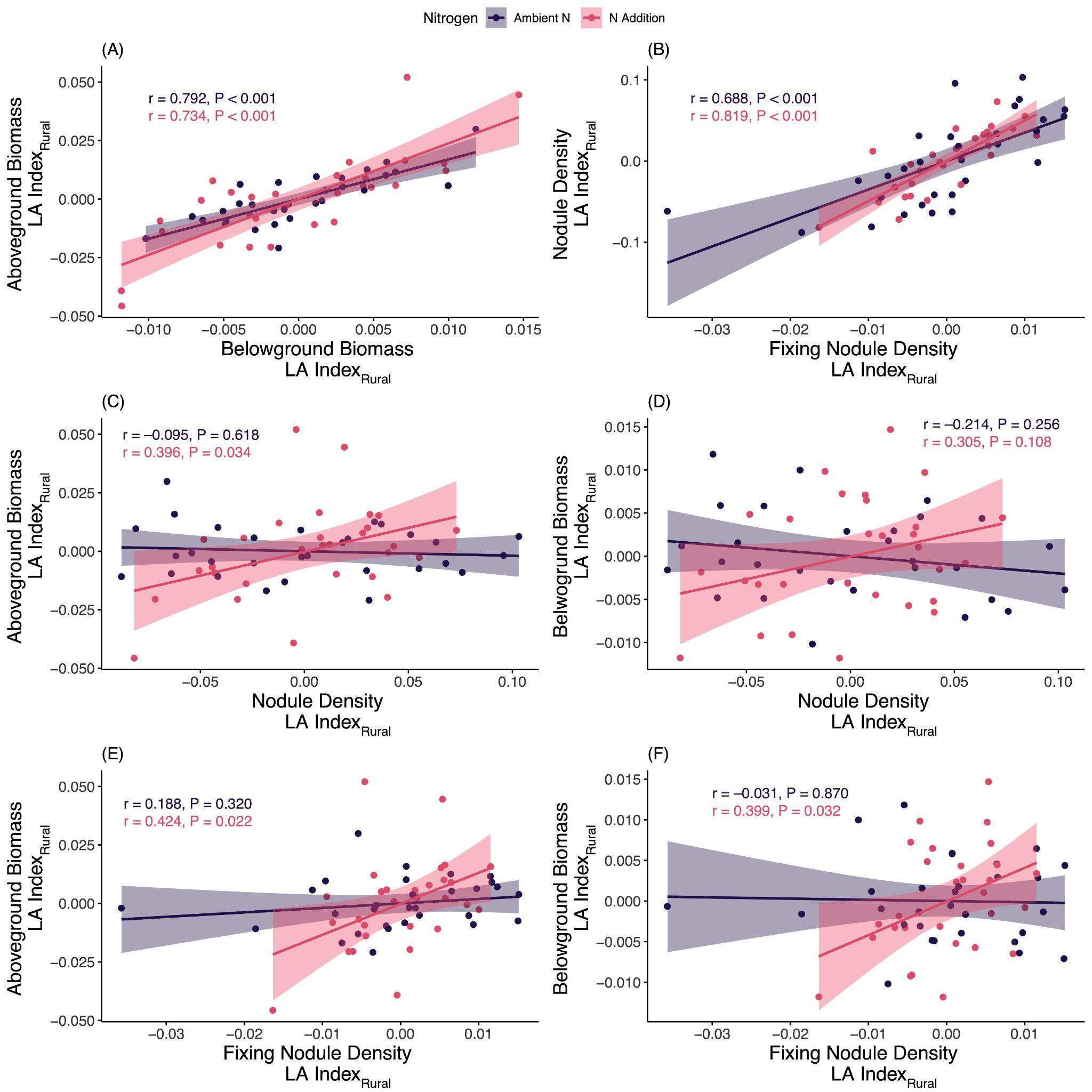

**Figure S2:** Plots showing all pairwise correlations between the local – nonlocal_Rural_ local adaptation index for aboveground biomass, belowground biomass, nodule density, and fixing nodule density. Separate correlations were calculated for each nitrogen treatment (ambient N = purple, N addition = pink). Lines represent the lines-of-best-fit (± 95% confidence interval). Inset text provides the correlation coefficient (r) and P-value for each correlation.

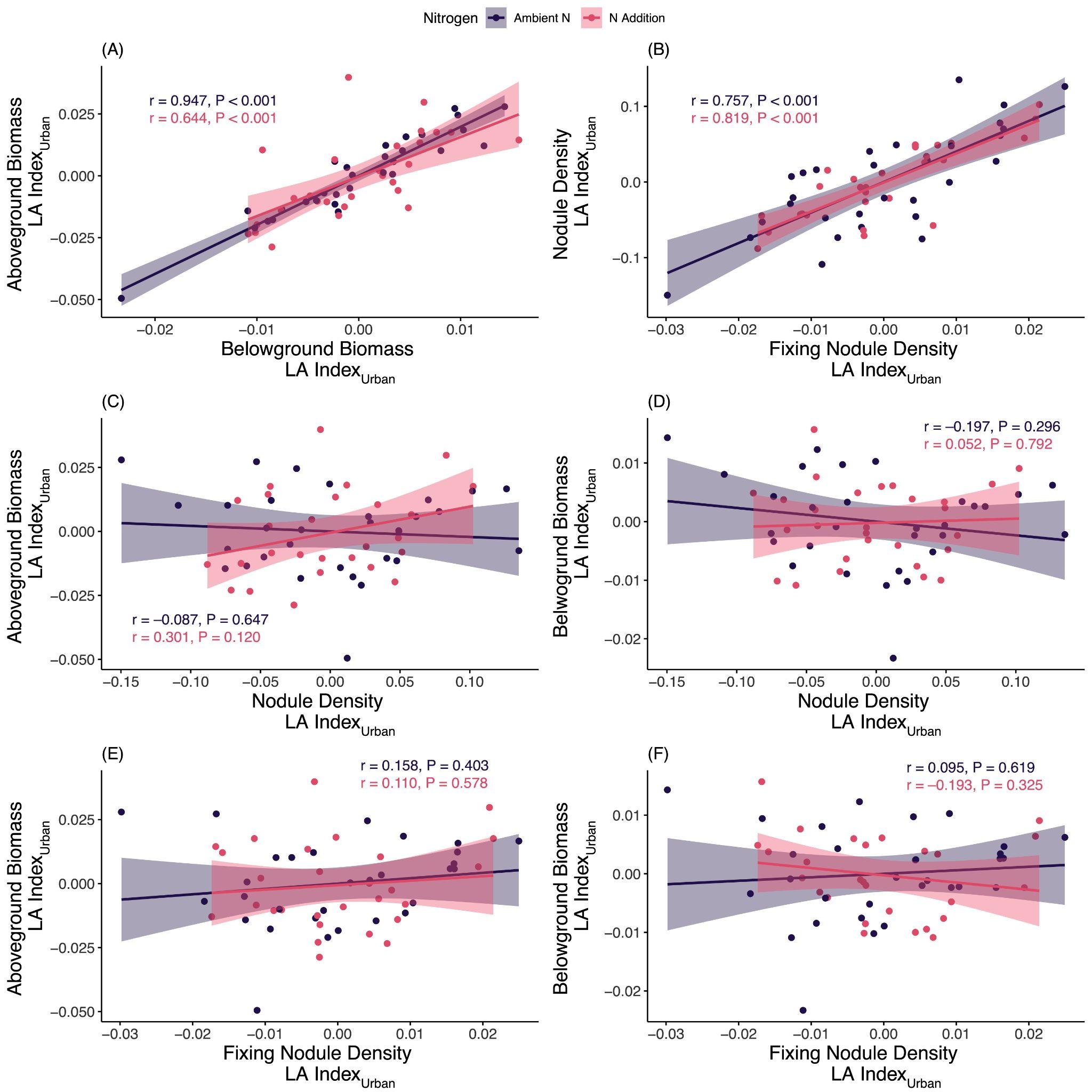

**Figure S3:** Plots showing all pairwise correlations between the local – nonlocal_Urban_ local adaptation index for aboveground biomass, belowground biomass, nodule density, and fixing nodule density. Separate correlations were calculated for each nitrogen treatment (ambient N = purple, N addition = pink). Lines represent the lines-of-best-fit (± 95% confidence interval). Inset text provides the correlation coefficient (r) and P-value for each correlation.

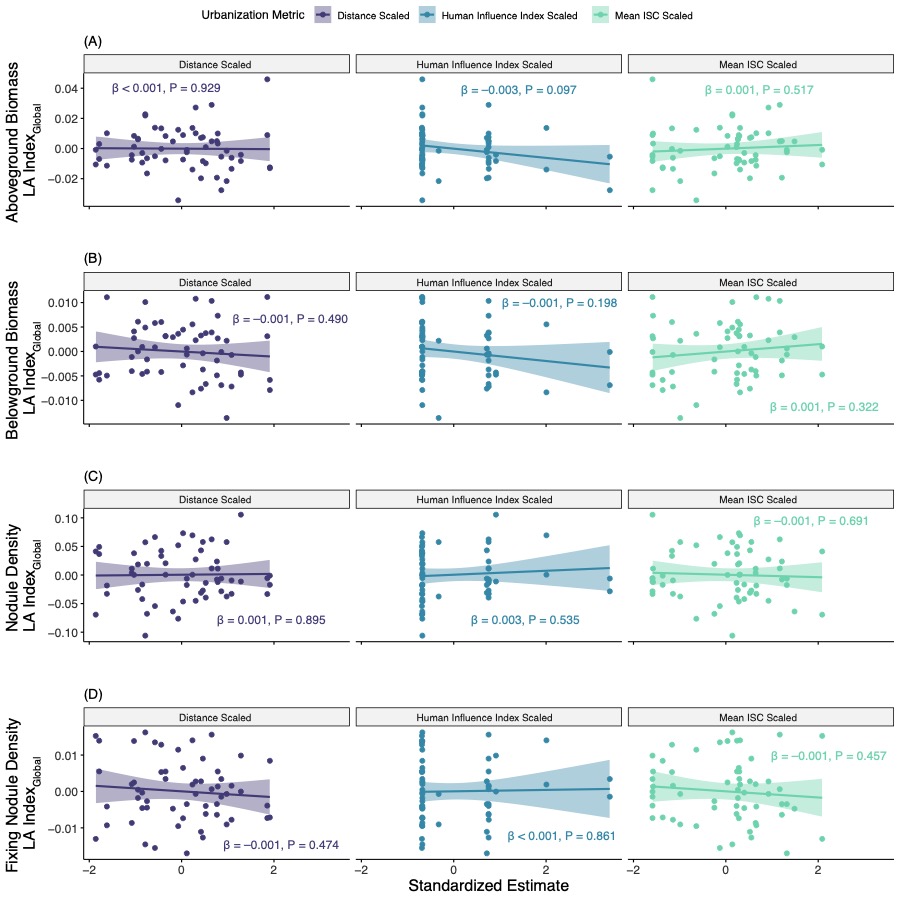

**Figure S4:** Facet plots of the relationship between the global local adaptation index (LA Index_Global_) for each of the fitness estimates [aboveground biomass (A) belowground biomass (B), nodule density (C), and fixing nodule density, (D)] and urbanization metrics [left column = distance from the city center, center column = human influence index, and right column = mean impervious surface cover (ISC)]. Lines represent the lines-of-best-fit (± 95% confidence interval). Inset text provides the slope parameter ($\beta$) and P-value for each linear regression.

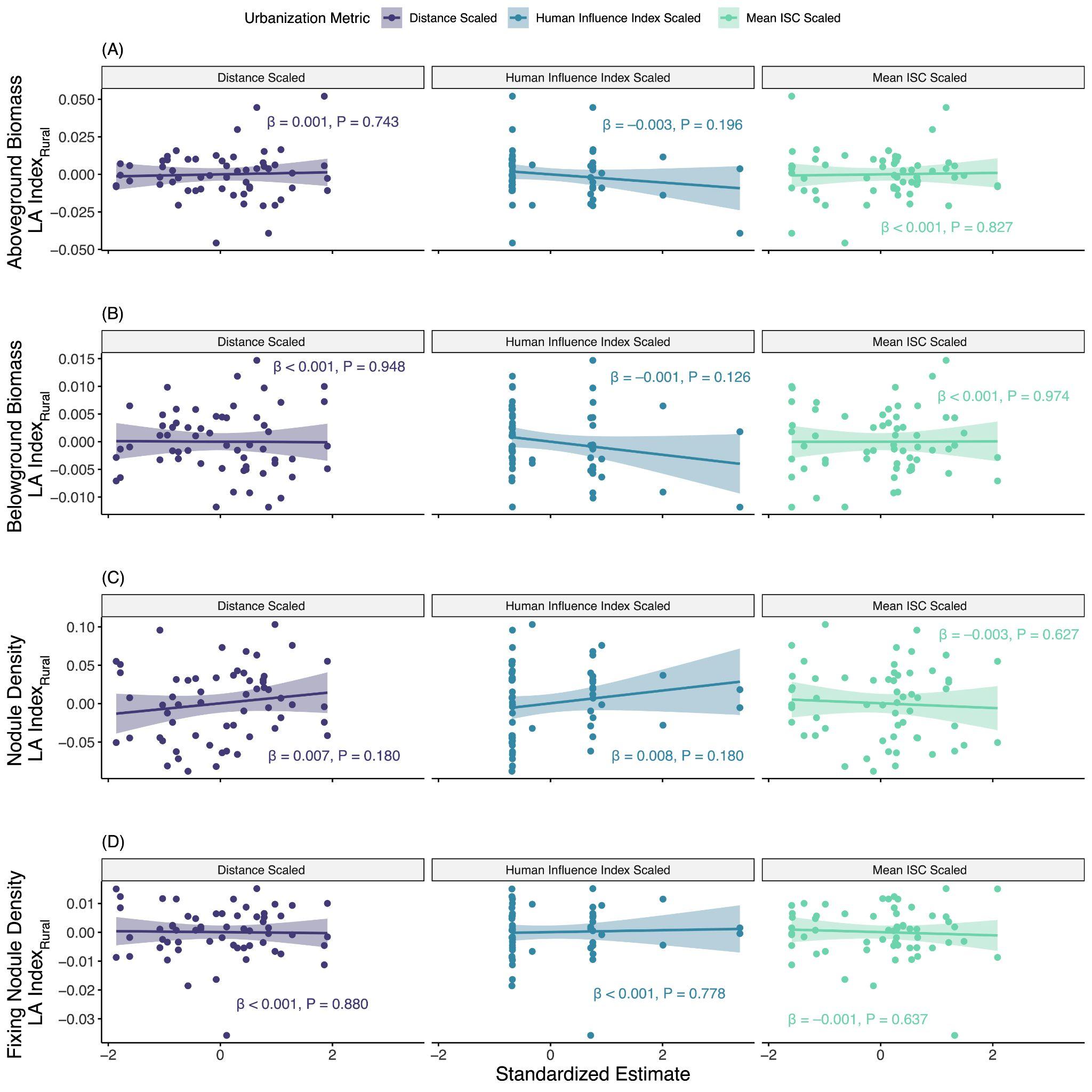

**Figure S5:** Facet plots of the relationship between the rural local adaptation index (LA Index_Rural_) for each of the fitness estimates [aboveground biomass (A) belowground biomass (B), nodule density (C), and fixing nodule density, (D)] and urbanization metrics [left column = distance from the city center, center column = human influence index, and right column = mean impervious surface cover (ISC)]. Lines represent the lines-of-best-fit (± 95% confidence interval). Inset text provides the slope parameter ($\beta$) and P-value for each linear regression.

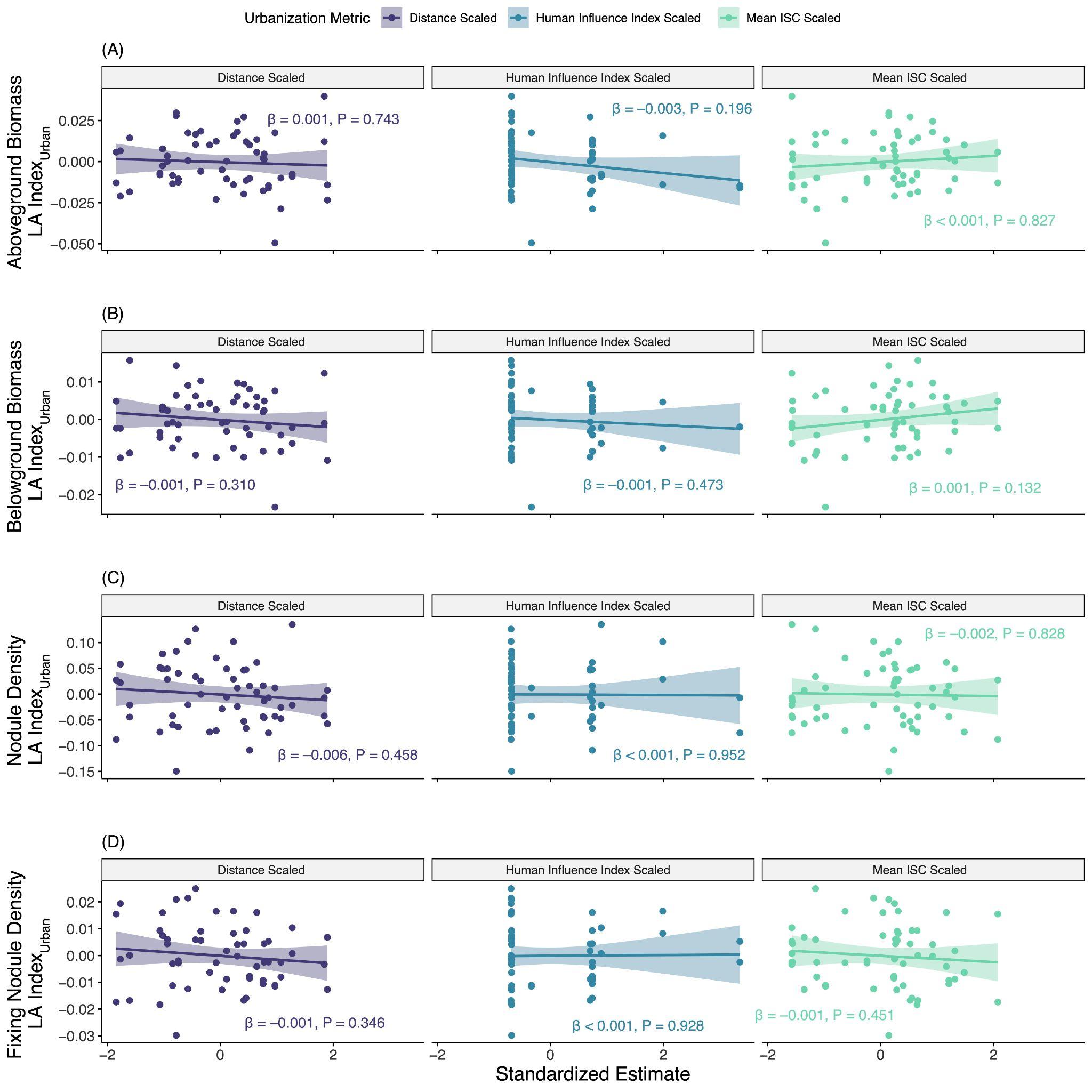

**Figure S6:** Facet plots of the relationship between the urban local adaptation index (LA Index_Urban_) for each of the fitness estimates [aboveground biomass (A) belowground biomass (B), nodule density (C), and fixing nodule density, (D)] and urbanization metrics [left column = distance from the city center, center column = human influence index, and right column = mean impervious surface cover (ISC)]. Lines represent the lines-of-best-fit (± 95% confidence interval). Inset text provides the slope parameter ($\beta$) and P-value for each linear regression.

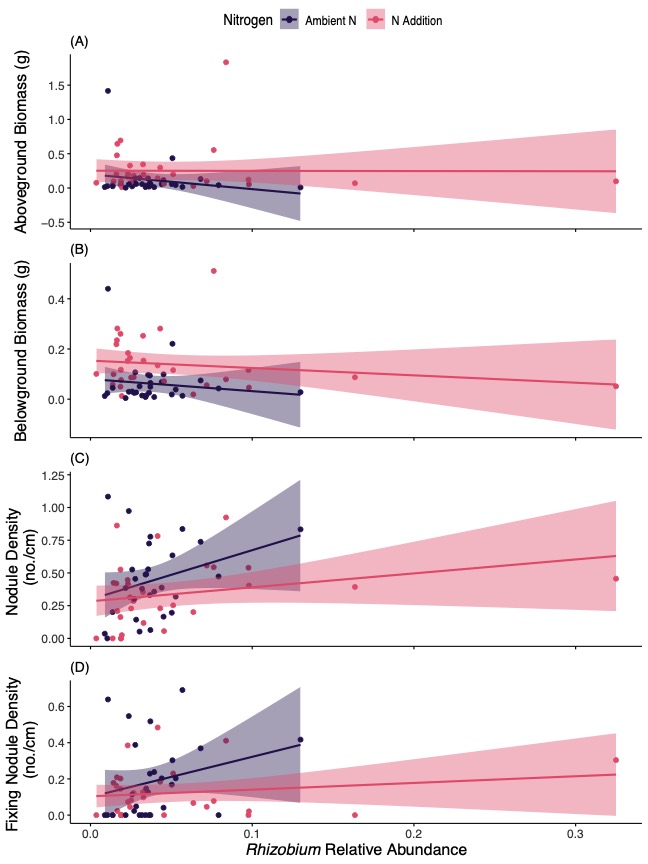

**Figure S7:** Plots of the relationship between fitness estimates and *Rhizobium* relative abundance for aboveground biomass (A), belowground biomass (B), nodule density (C), and fixing nodule density (D). Lines represent lines-of-best-fit (± 95% confidence interval) by nitrogen treatment (ambient N = purple, N addition = pink). Detailed test statistics are provided in Table S4.

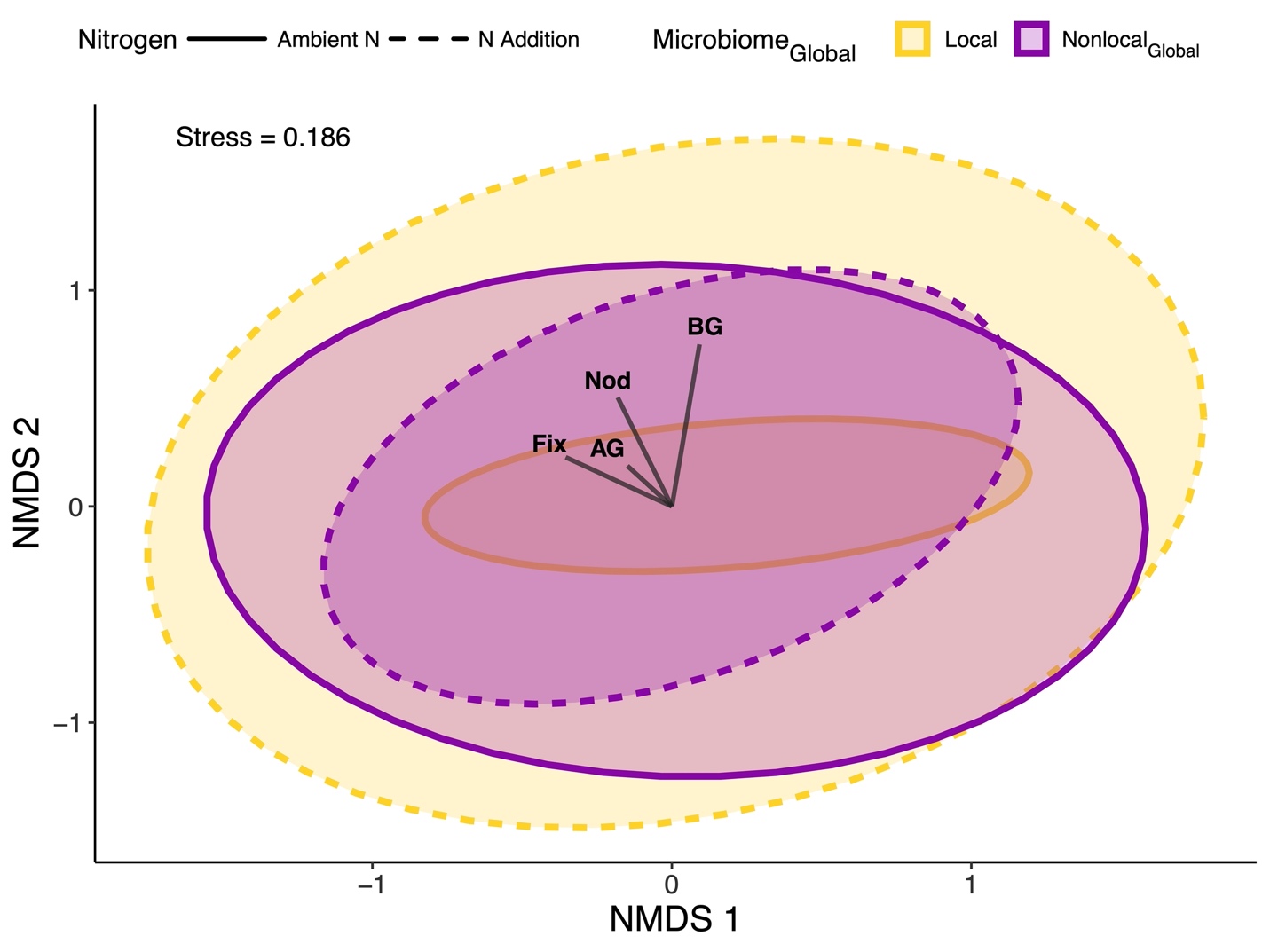

**Figure S8:** Non-metric multidimensional scaling plot (Bray-Curtis dissimilarity) of root microbial community composition by microbiome (local and nonlocal_Global_) and nitrogen (ambient N and N nddition) treatments. Lines indicate the relationship between community composition and the local adaptation index for each fitness estimate (aboveground biomass = AG, belowground biomass = BG, nodule density = Nod, and fixing nodule density = Fix). Detailed test statistics for the PERMANOVA are provided in Table S7.

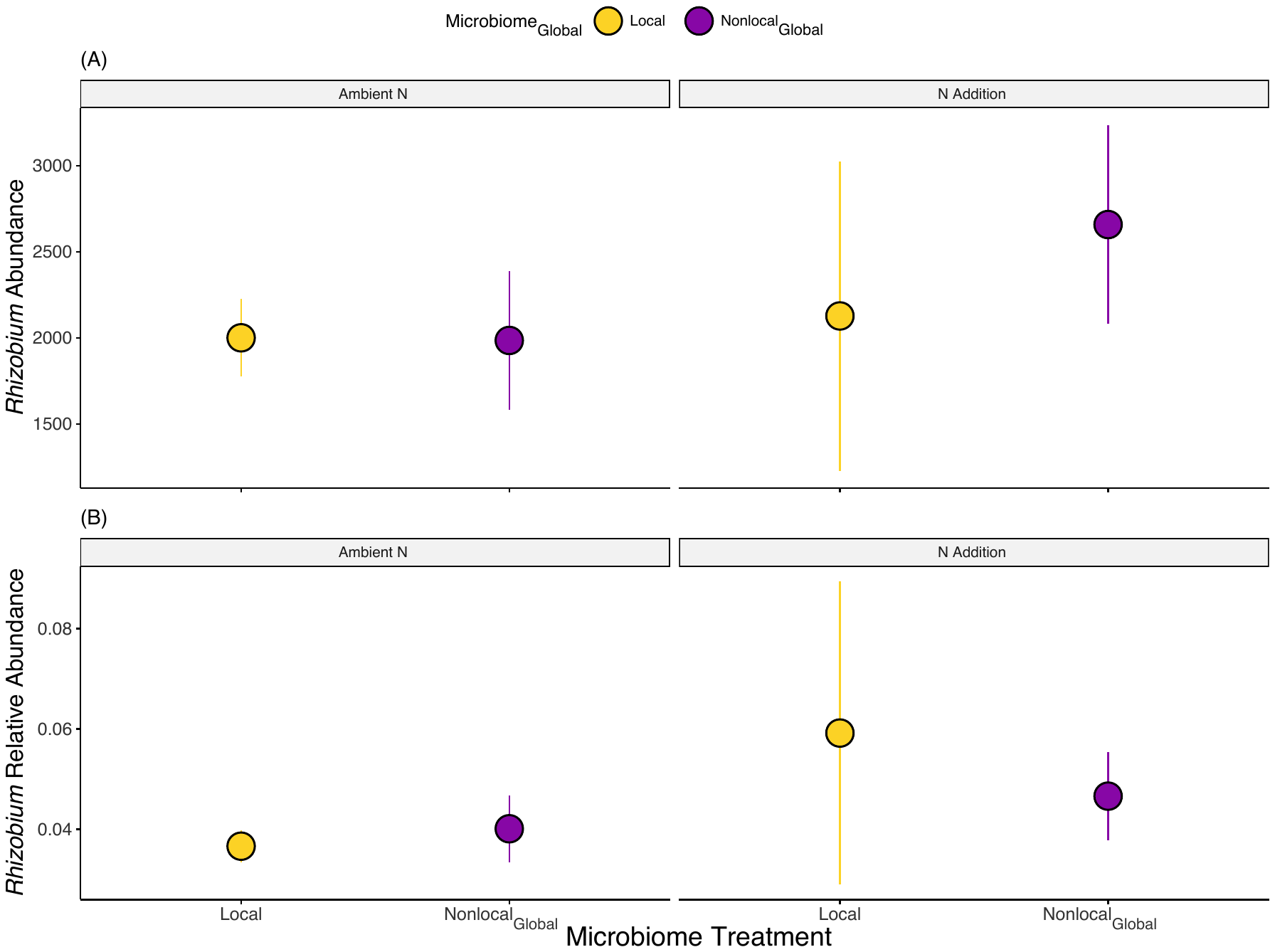

**Figure S9:** Plots of *Rhizobium* abundance (A) and relative abundance (B) by microbiome and nitrogen treatments. Points represent mean ± SE. Microbiome treatments are indicated as local (yellow) and nonlocal_Global_ (purple), and plots are faceted by nitrogen treatment (ambient N and N addition). Detailed test statistics are provided in Table S6.

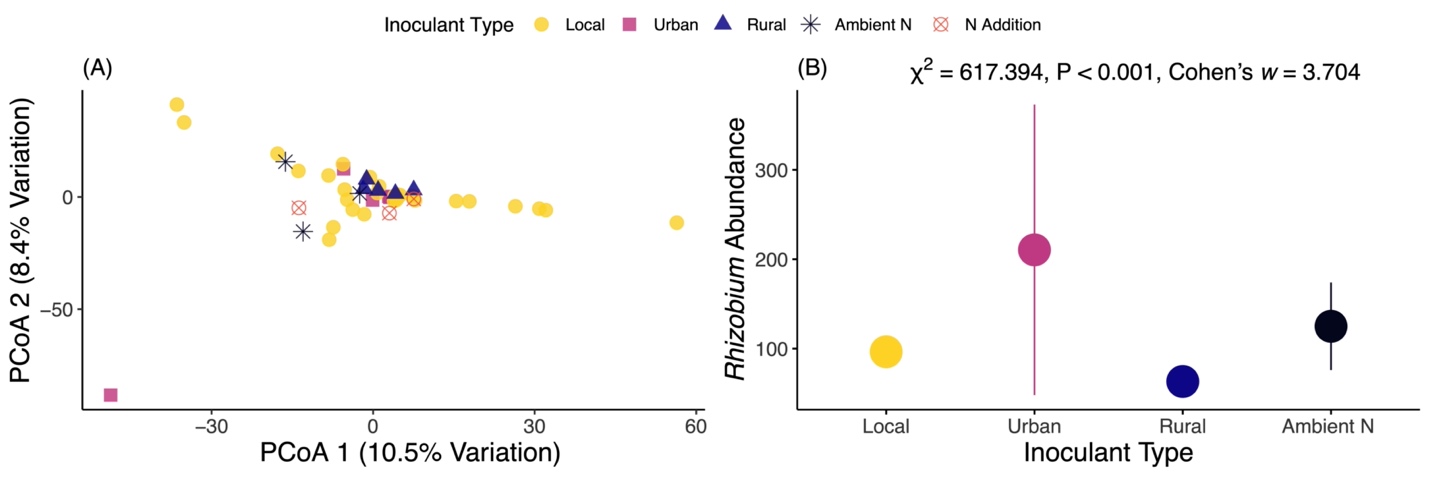

**Figure S10:** Principal coordinates analysis (PCoA) of microbiome inoculant communities (A). Local inoculants (yellow circles) were paired to each of the 30 focal populations, urban inoculants (pink squares) represent each of the 5 urban populations for the nonlocal_Urban_ inoculant, and rural inoculants (blue triangle) represent each of the 5 urban populations for the nonlocal_Rural_ inoculant. In contrast, ambient N (black stars) and N addition (orange square cross) were experimental controls that did not receive a live microbiome inoculant, only differing in their respective N fertilizer treatment. We also show the mean ± SE of *Rhizobium* abundance for each of the inoculants and experimental controls. Although there were overall differences in *Rhizobium* abundances by inoculant type (one-way ANOVA with Type III sums-of-squares χ^2^ = 617.394, P < 0.001, Cohen’s *w* = 3.704), the pairwise differences when *Rhizobium* were present were generally weak (range of Cohen’s *d* = 0.015-0.157). Note: no *Rhizobium* sequences were identified in the N addition experimental control and therefore those data are not presented.
